# Secreted dengue virus NS1 from infection is predominantly dimeric and in complex with high-density lipoprotein

**DOI:** 10.1101/2022.04.06.487425

**Authors:** Bing Liang Alvin Chew, An Qi Ngoh, Wint Wint Phoo, Kitti Wing Ki Chan, Zheng Ser, Nikhil K Tulsian, Shiao See Lim, Mei Jie Grace Weng, Satoru Watanabe, Milly M. Choy, Jenny G. Low, Eng Eong Ooi, Christiane Ruedl, Radoslaw M. Sobota, Subhash G. Vasudevan, Dahai Luo

## Abstract

Severe dengue infections are characterized by endothelial dysfunction shown to be associated with the secreted nonstructural protein 1 (sNS1), making it an attractive vaccine antigen and biotherapeutic target. To uncover the biologically relevant structure of sNS1, we obtained infection-derived sNS1 (isNS1) from DENV-infected Vero cells through immunoffinity purification instead of recombinant sNS1 (rsNS1) overexpressed in insect or mammalian cell lines. We found that isNS1 appeared as an approximately 250 kDa complex of NS1 and ApoA1 and further determined the cryoEM structures of isNS1 and its complex with a monoclonal antibody/Fab. Indeed, we found that the major species of isNS1 is a complex of the NS1 dimer partially embedded in a High-Density Lipoprotein (HDL) particle. Cross-linking mass spectrometry (XL-MS) studies confirmed that the isNS1 interacts with the major HDL component ApoA1 through interactions that map to the NS1 wing and hydrophobic domains. Furthermore, our studies demonstrated that the sNS1 in sera from DENV-infected mice and a human patient form a similar complex as isNS1. Our results report the molecular architecture of a biological form of sNS1 which may have implications for the molecular pathogenesis of dengue.

**Summary:** CryoEM structures of secreted dengue virus NS1 protein reveal dimers in complex with high-density lipoprotein.

## Introduction

Dengue virus (DENV) is a member of the flavivirus genus and is the cause of significant healthcare problems and economic burden worldwide, partially due to the lack of effective therapeutics and the limited efficacy of licensed vaccines, Dengvaxia® and QDENGA®^1,2^. While the majority of DENV infections are mild or asymptomatic, severe dengue infections, marked by vascular leakage, can be life-threatening and even fatal. The severity of dengue infections has been demonstrated to be correlated to high sera levels of sNS1 in clinical studies^3,4^. The viral nonstructural protein 1 (NS1) is a highly conserved and multifunctional protein that exists as intracellular membrane-associated NS1, a presumed GPI-anchored outer plasma membrane-associated NS1, and secreted NS1 (sNS1) during viral infection^5^. NS1 could interact with a myriad of proteins associated with its immune evasion and virotoxin roles^6^. The interactions of NS1 with viral E protein^7^ and the NS4A-2K-NS4B precursor protein^8^ have been shown to be important in virus production and replication respectively. Intracellular NS1 found in the ER lumen is an essential part of the membranous viral RNA replication compartment^7^. Upon release from the cells, sNS1 enters the blood circulation where *in vitro* and *in vivo* mouse evidence have demonstrated its ability to induce vascular permeability, a hallmark of severe dengue, either independently^9^ or through inducing pro-inflammatory responses^10^.

The high-resolution crystal structures of NS1 dimer have been reported for DENV^11^, WNV^11^, and ZIKV^12,13^ which provide a conserved molecular view of flaviviral NS1. Mature NS1 is 352 amino acids long with an apparent molecular mass between 40 – 50 kDa depending on its glycosylation state. NS1 has a three-domain architecture, a hydrophobic β-roll (residues 1-29), an α/β wing (38-151), and a β-ladder (181-352). The connector segments between the wing and β-ladder domains, residues 30-37 and 152-180, form a 3-stranded β-sheet. The dimer has a distinct cross shape with the wings extending from the central β-ladder which has an extended β-sheet that faces the hydrophobic β-roll and a ‘‘spaghetti loop” on the opposite hydrophilic outer face that lacks structured elements^11–13^. sNS1 was reported to be a barrel-shaped hexamer with lipid cargo held together by hydrophobic interactions based on biophysical and low-resolution EM analysis^14–16^. Recent cryoEM structures of recombinant sNS1 (rsNS1) from DENV-2 showed mainly tetrameric rsNS1 with only 3% of the population found to be in the hexameric form^17^. While earlier studies by Gutsche et al (2011) found that the lipid profile of DENV-1 infection-derived sNS1 to be similar to high-density lipoprotein (HDL)^15^, the same research group now showed that DENV-2 rsNS1 could dock onto HDL with direct visualization using negative stain EM analysis as reported by Benfrid et al. (2022)^18^. Additionally, sNS1 appears to utilize the scavenger receptor B1, a known receptor for HDL, as the cognate receptor in cultured cells^19^. Since the interactions of sNS1 with the target cells implicated in pathogenesis require the hydrophobic β-roll side to be exposed^11^, the mechanism by which the oligomeric sNS1 with its lipid load in the central channel dissociates and associates have been a topic of intense research interest^18,20,21^. However, the native structure of the extracellular NS1 during viral infection remains unclear.

To closely mimic of sNS1 circulating in dengue patients’ sera, we obtained infection-derived sNS1 (isNS1) from the culture supernatant of Vero cells infected with either the DENV2 WT (isNS1wt) or T164S mutant (isNS1ts). The T164S mutation in the greasy finger loop between the wing and β-ladder interdomain of NS1 was identified from a DENV2 epidemic in Cuba in 1997, where a correlation with enhanced clinical disease severity was observed^22^. We demonstrated that this single mutation in NS1 could directly cause lethality in mice and increased sNS1 secretion^23^, which served as an epidemiologically relevant tool to study the biochemical and structural characteristics of sNS1 in this study.

We determined that the isNS1 is a complex of the NS1 dimer embedded on a single HDL particle composed of ApoA1. By applying integrative structural tools, we report the cryoEM structural model of the purified isNS1:HDL complex at ∼8 Å resolution with protein-protein interactions between NS1 and ApoA1 defined by crosslinking mass spectrometry (XL-MS). Furthermore, the Ab56.2 binding site model in the cryoEM model of isNS1:ApoA1 complex was elucidated using hydrogen deuterium exchange mass spectrometry (HDX-MS) as well as NS1 peptide competition ELISA. Interrogation of DENV-infected mouse sera and a human patient suggests that the sNS1:HDL complex exists *in vivo* as a similar complex to isNS1.

## Results

### Native isNS1 is a complex of NS1 with ApoA1

We purified sNS1 using a monoclonal antibody 56.2 from the supernatant of the infected Vero cell cultures (Fig. 1a and Supplementary Fig. 1a-b). Initial negative stain screening of the isNS1wt (Supplementary Fig. 1c) showed some 2D classes of NS1 dimers presumably docked onto a spherical density, mirroring the *in vitro* reconstitution of crsNS1 with HDL complex negative stain microscopy results by Benfrid et al. (2022)^18^. Both immunoaffinity purified isNS1wt (Fig. 1b and Supplementary Fig. 2a) and isNS1ts (Supplementary Fig. 2a and 3b) retained a molecular size of ∼250 kDa as detected with Coomassie Blue (Fig. 1b and Supplementary Fig. 2a) and on a Western blot using Ab56.2 following separation on a Native-PAGE (Fig. 1b), as previously reported^23^. The isNS1wt migrated slower than the rsNS1 protein (a gift from Shu et al. 2022^17^), indicating possible conformational or oligomeric differences between the two NS1 species (Fig. 1b). On a reducing SDS-PAGE, we identified two major bands of approximately 50 kDa and 25 kDa in isNS1wt, the former corresponding to the single monomeric NS1 band seen in rsNS1 (Fig. 1c). Preliminary mass ID identified the 25 kDa protein as bovine apolipoprotein A1 (ApoA1), a major component of HDL in bovine serum-containing culture media, which has a reported molecular weight of 28 kDa^24^. Hence, the two bands were further validated on a Western blot, which confirmed the 50 kDa band as NS1 and the 25 kDa band as ApoA1 respectively (Fig. 1c). The same observations were seen in isNS1t (Supplementary Fig. 3).

**Fig. 1.**
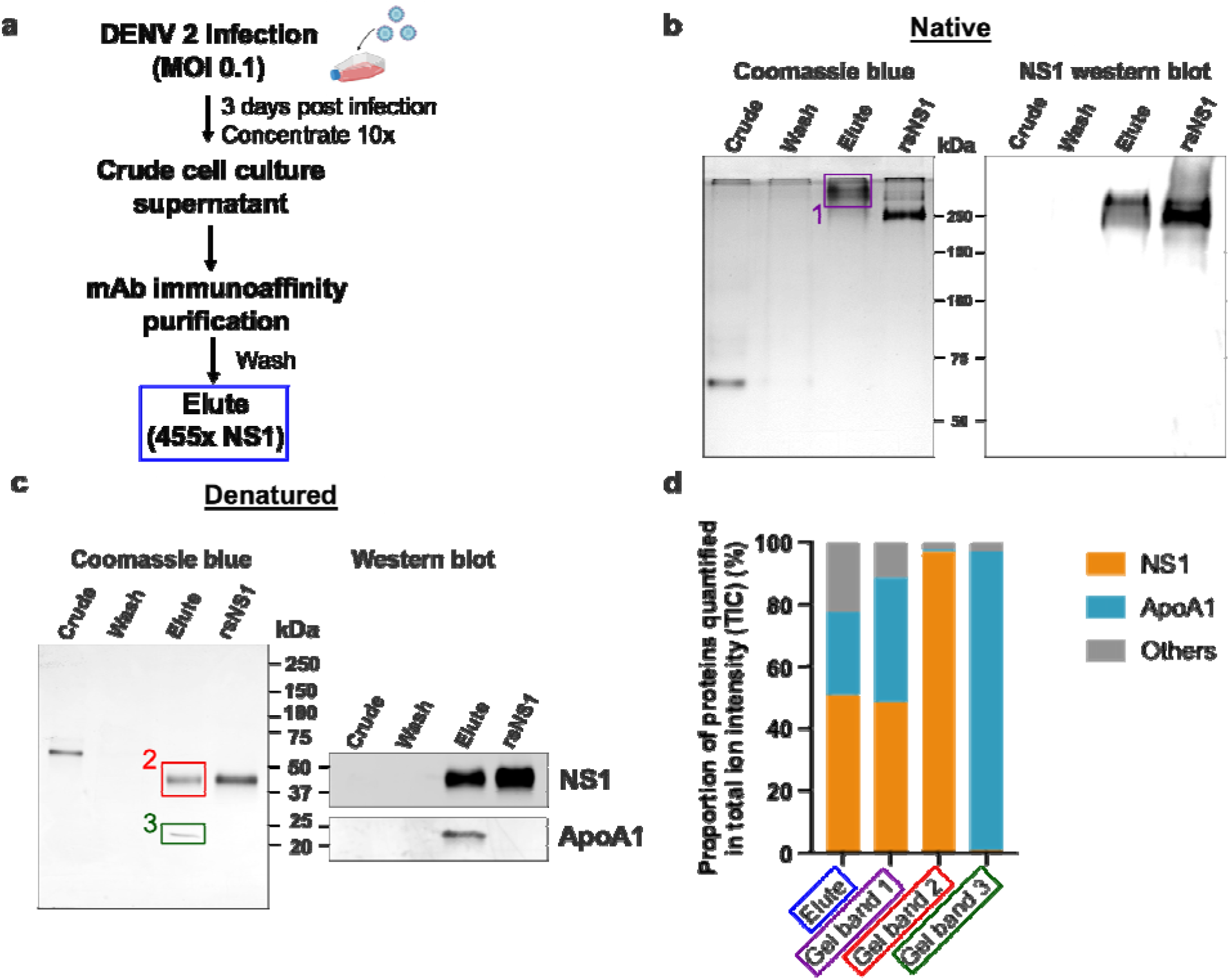
Composition of the secreted NS1 from DENV-infected Vero cells. DENV 2 WT cell culture supernatant was filtered, supplemented with protease inhibitor cocktail and 0.05% sodium azide, concentrated using a 100 kDa MWCO Vivaflow cassette and purified using 56.2 anti-NS1 antibody immunoaffinity chromatography. The eluted isNS1wt was dialysed against PBS, concentrated, and stored at –80°C until further use. **(a)** Schematic of isNS1 purification to illustrate the samples used for gel analyses. % NS1 is measured by the total amount of NS1 (quantified using the anti-NS1 ELISA kit (Bio-rad) as a percentage of total protein (quantified using the Bradford assay) found in each sample. Details of the % enrichment in NS1 along the purification process is as shown in Supplementary Fig. S1 B. **(b)** Coomassie blue detection of proteins from Crude, Wash and Elute immunoaffinity fractions for isNS1wt, with the recombinant sNS1 (rsNS1) obtained from Shu et al (2022)^17^ as a positive control, after separation on a 10% Native-PAGE gel (left). The Crude and Elute fractions contain 1 µg of total protein. The Wash fraction contains approximately 100 ng of total protein in maximum well volume of the gel. The same set of samples were also subjected to a western blot detection of NS1 using 56.2 anti-NS1 antibody after separation on a 10% Native-PAGE (right). The Crude and Elute fractions contain 500 ng of total protein. The Wash fraction contains approximately 100 ng of total protein in maximum well volume of the gel. **(c)** Coomassie blue detection of proteins from Crude, Wash and Elute immunoaffinity fractions for isNS1wt and rsNS1^17^, after separation on a 4-20% reducing SDS-PAGE gel. The Crude and Elute fractions contain 1 µg of total protein. The Wash fraction contains approximately 100 ng of total protein in maximum well volume of the gel. Similarly, the same set of samples were also subjected to a western blot detection of NS1 and ApoA1 using 56.2 anti-NS1 antibody or ApoA1 antibody (Biorbyt, orb10643) respectively, after separation on a 4-20% reducing SDS-PAGE (right). **(d)** In-gel protein identification of the purified isNS1wt by liquid chromatography mass spectrometry (LC-MS). Proportion of NS1, ApoA1 and other unidentified proteins quantified in total ion intensity, obtained from the following samples: Elute in solution (boxed in blue), 250 kDa gel band (boxed in purple), 50 kDa gel band (boxed in red) and 25 kDa gel band (green). The boxed gel bands are from representative gels showing the different protein species found while the actual gel bands used for protein identification by LC-MS are as shown in Supplementary Fig. 2a-b.

Next, we elucidated the composition of the prominent 250 kDa band observed in the Native-PAGE by label-free quantification (LFQ) analysis of proteins using liquid chromatography–mass spectrometry (LC-MS). We obtained LC-MS data for the ∼250 kDa protein excised from the Native-PAGE separation of the immunoaffinity purified isNS1wt (represented in Fig. 1b, purple box as Gel Band 1); an aliquot of the elute fraction from immunoaffinity purification (depicted in Fig. 1a blue box and referred to as “Elute”) and the 50 kDa and 25 kDa bands excised after separation of the elute fraction on a reducing SDS-PAGE (Fig. 1c, represented by red and green boxed labelled as Gel Band 2 and 3 respectively) as positive controls. The raw data for the ion intensities of the samples tested are as detailed in Supplementary File 1. We found that the combined LFQ intensity of NS1 and ApoA1 accounted for more than 77.6% and 88.6% of the total ion intensity count (TIC) detected from the elute fraction (Fig. 1a, blue box) and the 250 kDa band (Fig. 1b, the purple box labelled ‘1’) respectively, indicating that NS1 and ApoA1 were the major proteins in the elute fraction and the ∼250 kDa band from immunoaffinity purified isNS1wt (Fig. 1d). Accordingly, as depicted in Fig. 1d, NS1 accounted for 96.85% of the TIC for the 50 kDa band (Fig. 1c, red box ‘2’), while the intensity of ApoA1 accounted for 96.23% of the TIC for the 25 kDa band (Fig. 1c, green box ‘3’). Similar LFQ analysis of LC-MS data for the purified isNS1ts mutant showed that NS1 and ApoA1 were the major components (Supplementary Fig. 2) of the respective Elute and ∼250 kDa gel band on Native-PAGE (Supplementary Fig. 2a; 3a, purple box). The corresponding 50 kDa and 25 kDa gel bands on the reducing SDS-PAGE (Supplementary Fig. 2b, 3b) identified as NS1 and ApoA1 by immunoblotting with the respective antibodies (Supplementary Fig. 3b), were further confirmed by LC-MS as NS1 and ApoA1 (Supplementary Fig. 2c, 3c, Supplementary file 1). Taken together, the results from the Native PAGE and LFQ by LC-MS strongly indicated that the isNS1 purified from the supernatant of infected cells *in vitro* is an approximately 250 kDa complex of NS1 and ApoA1.

### CryoEM structure of isNS1

To gain insights into the molecular organization of this native isNS1, we determined the cryoEM structure of isNS1 (Supplementary Table 1 and Supplementary Fig. 4-7). The 2D classes of isNS1ts are observed to be heterogenous spheres with some internal features, some of which resembling the cross-shaped NS1 protruding from the sphere (Supplementary Fig. 4b). However, the 3D reconstruction of isNS1ts alone was observed to have no discernible features beyond a spherical-like density with varying dimensions averaging around 106 Å by 77 Å (Supplementary Fig. 4b-c). To obtain higher resolution structural information on the complex, we collected data for the ternary complexes of the isNS1wt:Ab56.2 isNS1wt:Fab56.2 (Fig. 2a and Supplementary Fig. 5-6). The use of antibodies as fiduciary markers is a well-proven approach for solving the structures of small proteins^25,26^. Overall, both samples resulted in similarly distinguishable 2D class averages (Fig. 2b-c) which show asymmetrical binding of one unit of Fab56.2 to the dimeric cross-shaped sNS1 associated with a lower-density sphere, presumably HDL.

**Fig. 2.**
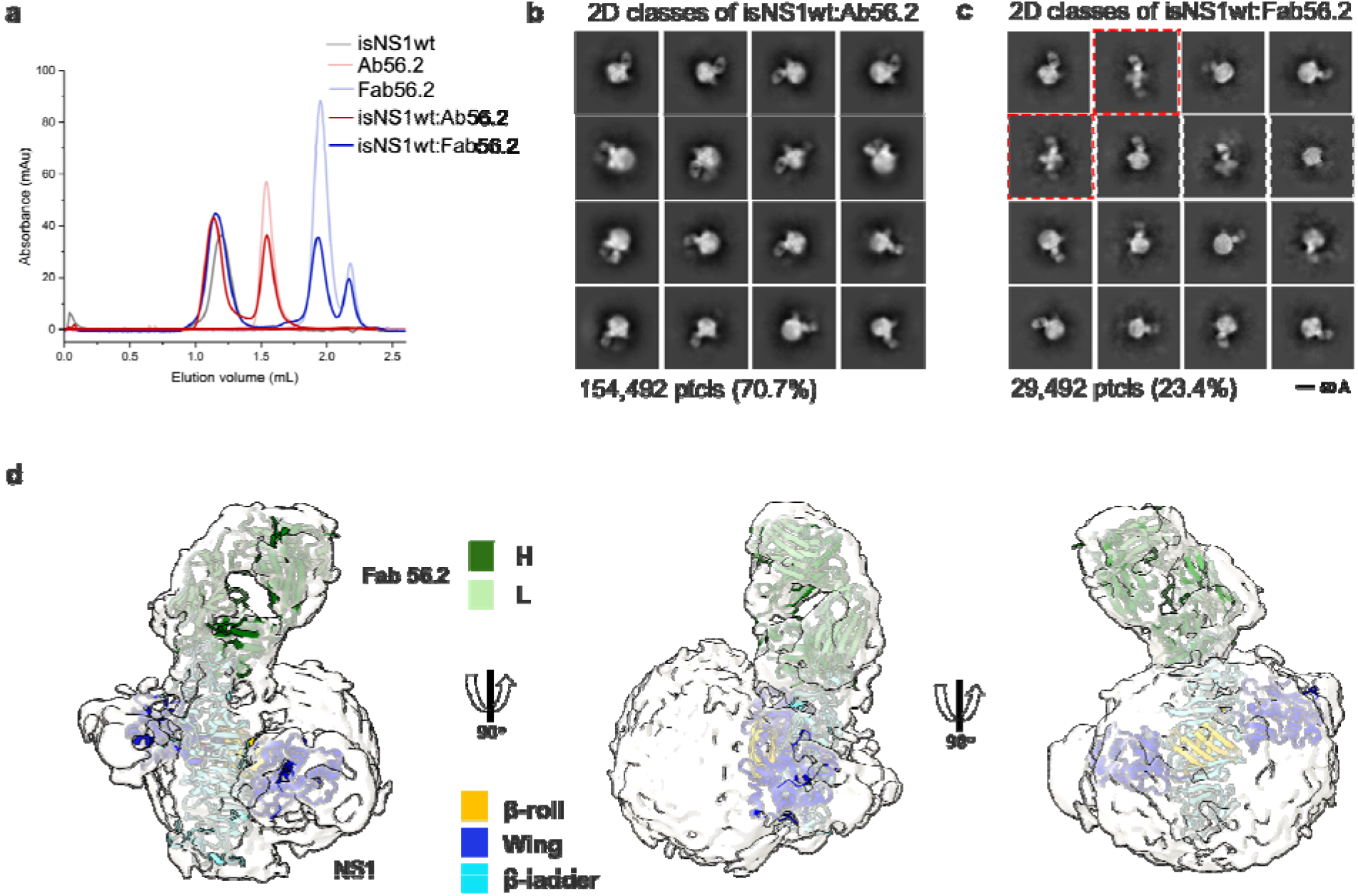
CryoEM analysis of secreted NS1 in complex with antibody Ab56.2. **(a)** Size exclusion chromatography was run on a Superdex 200 increase 3.2/300 GL column connected to the ÄKTA purifier with a flow rate of 0.075 mL/min in PBS (pH 7.4) for purified isNS1wt (gray), Ab56.2 (faded red), Fab56.2 (faded blue), isNS1wt:Ab56.2 (red) and isNS1wt:Fab56.2 (blue). A slight leftward shift in elution volume was observed for isNS1wt upon complexing with Ab56.2 and Fab56.2. 2D class averages of **(b)** isNS1wt:Ab56.2 and **(c)** isNS1wt:Fab56.2 showing representative sub-class of the Fab56.2:isNS1:HDL particles with black scale bar, 50 Å. The corresponding number of particles and percentages are listed below the respective boxes. Red dashed line boxes highlight two rare views consisting of 1033 particles (3.5%) only seen in isNS1wt:Fab56.2 sample. **(d)** Model of isNS1wt:Fab56.2 predicted structures rigid body fitted in the CryoEM map of Fab56.2:isNS1:HDL (grey, contoured at 0.14) with correlation value of 0.75 to the fitted regions (map simulated from atoms at 5 Å). isNS1wt is coloured by its three domains, namely the β-roll (orange), wing (blue), and β-ladder (cyan). Fab56.2 is coloured by it heavy chain (dark green) and light chain (light green).

In the isNS1wt:Fab56.2 ternary complex dataset, we further observed two apparent but rare sub-classes of free sNS1wt dimer bound to two units of Fab56.2 representing 3.5% of the total population of picked particles (Fig. 2c, red dashed boxes). While the cryoEM map reconstructions remain limited at low resolutions (Supplementary Fig. 5-6), they could be fitted with the NS1 dimer and Fab56.2 models (Fig. 2d) predicted separately using AlphaFold2^27,28^, with a correlation value of 0.75 to the fitted regions using a simulated 5 Å map from the NS1 dimer and Fab models. The AlphaFold2 predicted model of the NS1 dimer was used despite the availability of the crystallographic dimer model (PBS ID: 6WER)^29^ given their high similarity (RMSD: 0.67) and that the latter had a few unmodelled residues (aa 118-128). The map-model fitting showed that the epitope sites were at the tip of the C-terminal β-ladder region. The approximate dimensions of the spherical HDL could also be measured at 82 Å by 65 Å (Supplementary Fig. 5-6) which aligned with existing reports of spherical HDL being measured at 70-120 Å in diameter with predominantly homodimeric ApoA1 arranged as an anti-parallel double-belt stabilized by intermolecular salt bridges^30,31^. We predicted a model of an antiparallel dimer of ApoA1 using AlphaFold2^27,28^ without the first 58 residues of ApoA1 as the N-terminal domain is highly flexible^30^. The model has a cross-sectional distance of approximately 78 Å and fits into the spherical density in the cryoEM map. The NS1 dimer appears semi-embedded on the HDL particle through its hydrophobic surface, with a conformation that exposed only one end of the β-ladder domain to which Ab56.2 and Fab56.2 bind. Overall, the cryoEM model indicated that the purified ∼250 kDa isNS1 is a complex of a NS1 dimer and HDL, consistent with the gel and LFQ analysis of LC-MS data presented above.

Similarly, we determined the structures of the ternary complex of isNS1ts mutant with Fab56.2 (Fig. 3a). Interestingly, 2D class averages of isNS1ts:Fab56.2 dataset showed a significant sub-population of a free NS1ts dimer bound to two units of Fab56.2 (53.7%), in addition to the HDL spheres (23.6%) and isNS1ts dimer:HDL:Fab56.2 (22.7%) classes (Fig. 3b and Supplementary Fig. 7). The predicted structure of the isNS1ts dimer bound to Fab56.2 could be fitted with an overall correlation value of 0.5 using map simulated from atoms at 5 Å (Fig. 3c). Achieving atomic resolution to map the Fab binding interface with the NS1 β-ladder domain remains a challenge due to the preferred orientation (Supplementary Fig. 7). A comparison of the cryoEM density maps of free isNS1ts dimer:Fab56.2 (Fig. 3D, grey) and isNS1ts dimer:HDL:Fab56.2 ternary complex (Fig. 3d, yellow) fitted with a correlation value of 0.71 based on the NS1 map region, revealed a pose difference of the Fab56.2 between the free isNS1ts dimer and the HDL-bound isNS1ts dimer (Fig. 3d, inset). Further comparison of the density maps of the isNS1 dimer:HDL population in the Fab56.2 ternary complexes of isNS1wt (Fig. 3e, purple) and isNS1ts (Fig. 3e, yellow) revealed conformational similarities of the isNS1 dimer:HDL complex.

**Fig. 3.**
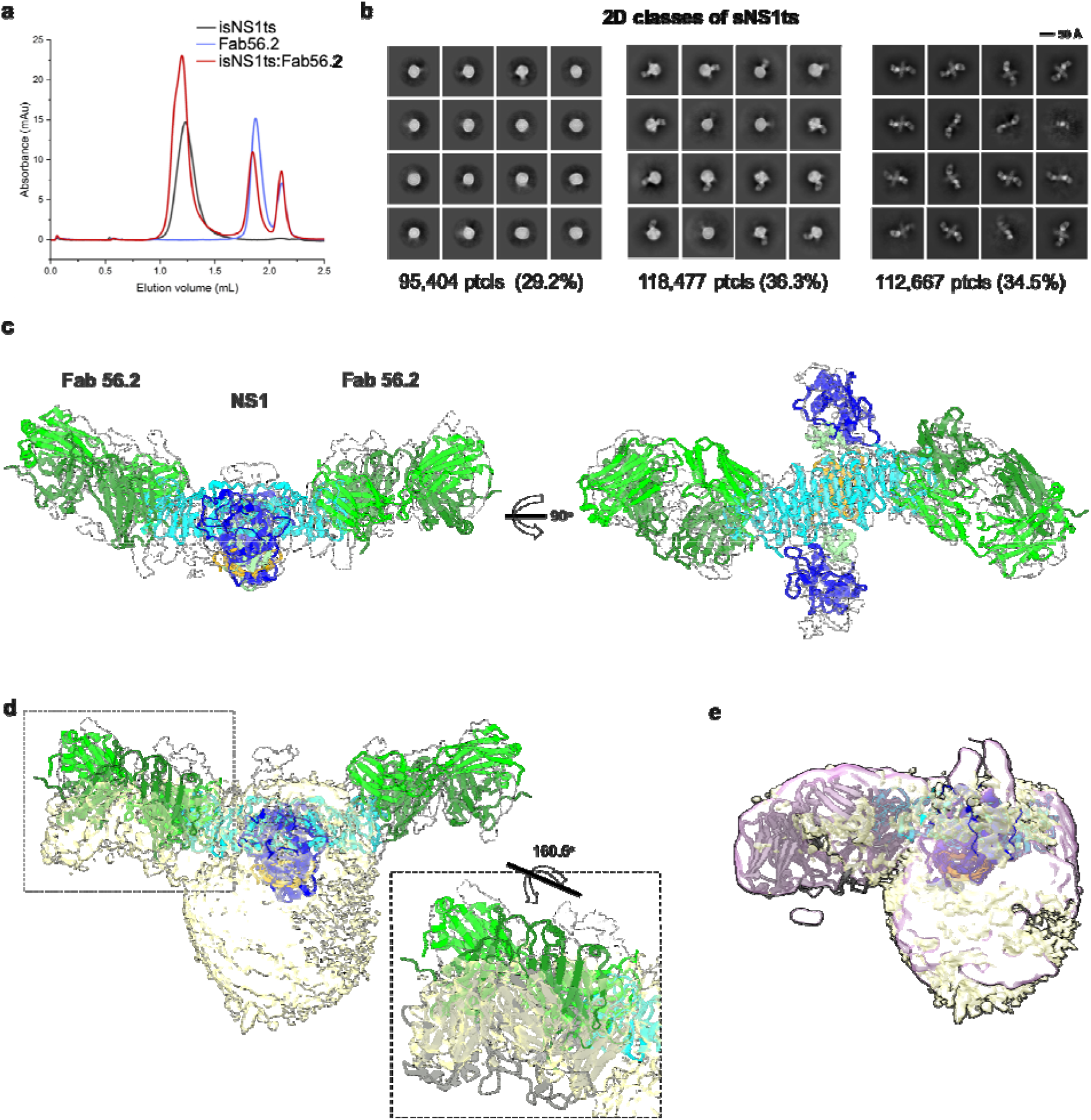
Secreted NS1 forms free dimers in complex with antibody Ab56.2. **(a)** Size exclusion chromatography was run on a Superdex 200 increase 3.2/300 GL column connected to th AKTA purifier with a flow rate of 0.075mL/min in PBS (pH 7.4) for purified isNS1ts (gray), Fab56.2 (faded blue) and isNS1ts:Fab56.2 (red). A similar leftward shift in elution volume wa also observed for isNS1ts upon complexing with Fab56.2. **(b)** 2D class averages of isNS1ts:Fab56.2 dataset with 326,548 particles picked and separated into 3 distinct populations, HDL spheres, Fab56.2:isNS1:HDL, and free isNS1:Fab56.2. Black scale bar, 50 Å, as indicated. The corresponding number of particles and percentages are listed below the respective boxes. **(c)** Model of isNS1ts dimer and Fab56.2 predicted structures rigid body fitted in the isNS1ts:Fab56.2 density map (grey, contoured at 0.14) with correlation value of 0.53 (overall, map simulated from atoms at 5 Å). isNS1ts is coloured by its three domains, namely the β-roll (orange), wing (blue), and β-ladder (cyan). Fab56.2 is coloured by its heavy chain (dark green) and light chain (light green). **(d)** Density map fitting between isNS1ts:Fab56.2 (grey, contoured at 0.14) to Fab56.2:NS1ts:HDL (yellow, contoured at 0.1) with correlation value of 0.53 (overall) and 0.7137 (on D2NS1 map region only). Inset shows the difference in the pose of Fab from the free NS1 form to the HDL-bound form. (e) Density map fitting between Fab56.2:isNS1wt:HDL (purple, contoured at 0.05) to Fab56.2:NS1ts:HDL (yellow) with correlation value of 0.72 (overall).

### XL-MS maps the inter- and intra-molecular architecture of the isNS1

Next to refine our understanding of the NS1 dimer:HDL complex of the isNS1 indicated by the LFQ of LC-MS (Fig. 1d and Supplementary Fig. 3d) and cryoEM results (Fig. 2d and 3d), we probed the interactions between NS1 and ApoA1 - the major protein moiety of HDL - using crosslinking mass spectrometry (XL-MS)^32^ on the immunoaffinity purification eluate fraction (Fig. 4a, Supplementary Fig. 8a-b). We identified 28 NS1:NS1 and 29 ApoA1:ApoA1 intramolecular crosslinks, as well as 25 NS1:ApoA1 intermolecular crosslinks for isNS1wt (Fig. 4b; Supplementary File 2a-b). As depicted in Fig. 4b-c, the NS1 residues that are involved in both intra- and inter-molecular crosslinks were located mainly on the β-roll and wing domains, with ∼34% of the NS1:ApoA1 crosslinks being located on the β-roll domain. The NS1:NS1 intramolecular crosslink pairs were further validated on the predicted NS1 dimer model by implementing a 30 Å cut-off distance for the Cα atoms of crosslinked residues, a criterion that was met by 85% of the NS1:NS1 intramolecular crosslinks (24 out of 28; Fig. 4d). The NS1:ApoA1 intermolecular crosslinks for the immunoaffinity purified isNS1wt were found to be located between the β-roll and wing domains of NS1 and the helices 3, 4, 8 and 10 of ApoA1 (Fig. 4b, d). Applying the same cut-off value of <30 Å distance between the Cα atoms of the NS1:ApoA1intermolecular crosslinked residues and validating it on the cryoEM model of isNS1wt in this study (Fig. 4d-e) satisfied 76% of the crosslinks (19 out of 25 intermolecular crosslinks between NS1 dimer and ApoA1). In the case of ApoA1:ApoA1 intramolecular crosslinks, the <30 Å distance between their Cα atoms cut-off satisfied 62% of the crosslinks (18 out of 29 ApoA1:ApoA1 intramolecular crosslinks). The higher number of violated crosslinks in ApoA1:ApoA1 compared to those in NS1:NS1 could be due to the flexible and dynamic nature of ApoA1 dimers which may prevent it from assuming a fixed relative position among the ApoA1 dimer population in the HDL. Overall, the XL-MS data and structural validation agreed with the cryo-EM model of isNS1 being a complex of the NS1 dimer embedded on a HDL particle composed of an ApoA1 dimer (Fig. 4d-e).

**Fig. 4.**
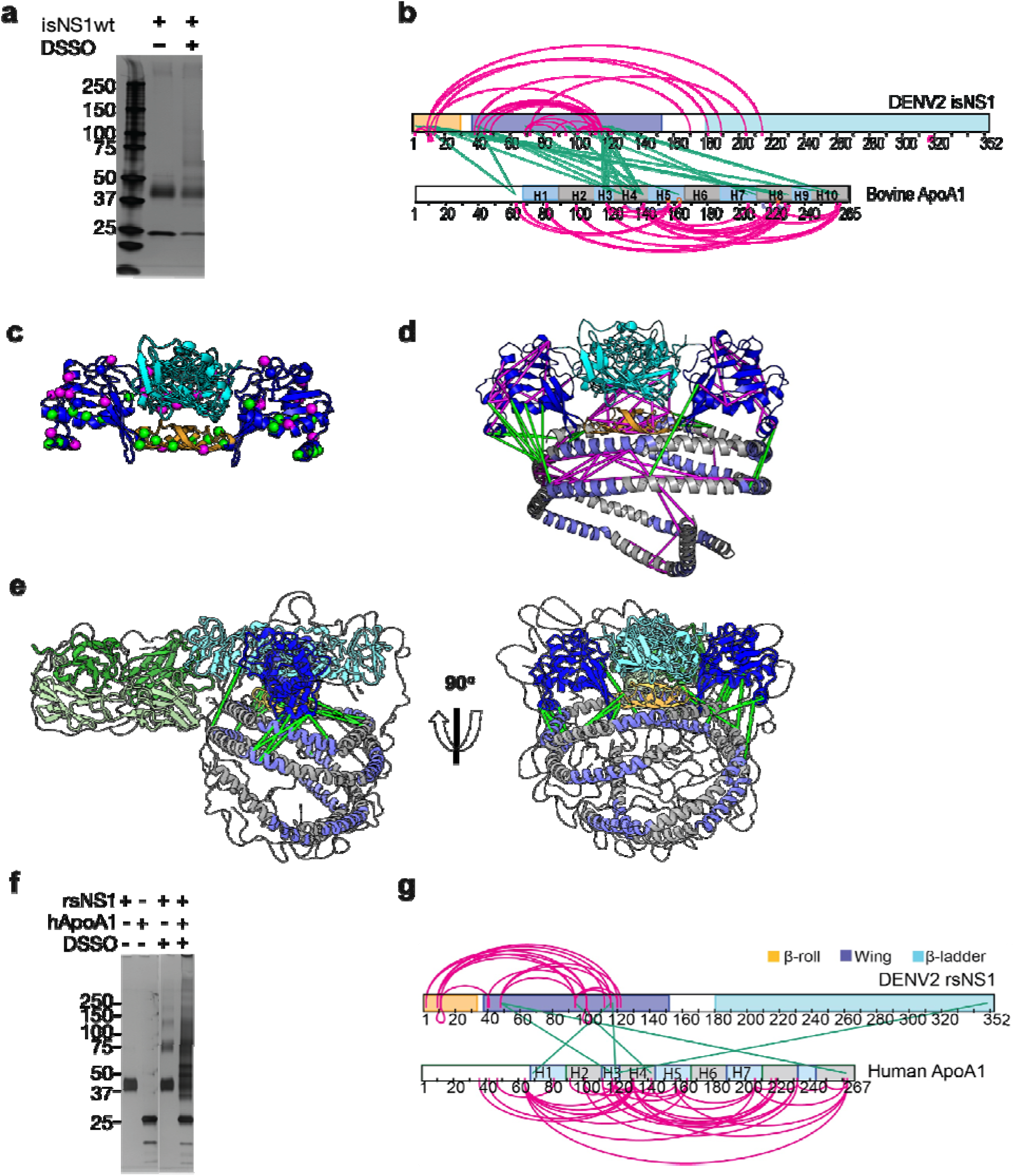
Interaction sites of NS1:ApoA1 complex identification by crosslinking mas spectrometry. **(a)** SDS-PAGE analysis of isNS1wt with or without the addition of DSSO crosslinker. **(b)** The identified crosslinks are visualised on the NS1 and bovine ApoA1 constructs. The intramolecular (NS1:NS1 and ApoA1:ApoA1) crosslinks are in magenta. Th intermolecular NS1:ApoA1 crosslinks are in green. **(c)** The isNS1 residues that are involved in NS1:NS1 interactions are visualised on the NS1 dimer model. The NS1 cartoon model i coloured by its three domains, namely the β-roll (orange), wing (blue), and β-ladder (cyan) with the intramolecular (magenta) and intermolecular (green) crosslinking sites depicted as spheres. **(d)** The overall model interpretation of NS1:ApoA1 complex within the crosslinker theoretical distance cut-off at =< 30 Å as depicted. ApoA1 dimer cartoon model with its conserved helices as labelled coloured in intervals of grey and light purple. **(e)** The NS1:ApoA1 dimer model with validated crosslinks were fitted into the cryoEM envelope. **(f)** SDS-PAGE analysis of crosslinked rsNS1 alone or with human HDL **(Lanes 3-5)**. Non-crosslinked rsNS1 and Human HDL are the control **(Lanes 1-2)**. The crosslinked rsNS1 can be seen in higher oligomers **(Lane 3)**. **(g)** Identified crosslinks are mapped on the NS1 and ApoA1 constructs, coloured as per panel B.

Additionally, the XL-MS mapping of isNS1ts mutant (Supplementary Fig. 8c-d) followed by rigid-body fitting into the corresponding cryoEM map (Supplementary Fig. 8e) revealed a similar crosslink pattern to isNS1wt, except that there were fewer crosslink sites (Supplementary Fig. 8f, Supplementary File. 2a). This indicated that isNS1ts dimer may have a weaker affinity to HDL in comparison to the isNS1wt dimer. This finding is consistent with the cryoEM data for isNS1ts where we found that free isNS1ts dimers bound to Fab56.2 formed a significant sub-population (34.5%) of the total picked particles (Fig. 3b, Supplementary Fig. 7b).

Given the reported structures of hexameric and tetrameric rsNS1 from mammalian expression systems, insect cells^11,15,16,29^ and Expi293 HEK cells^17^, we further examined whether rsNS1 oligomerizes in the presence of exogenously added human HDL, using XL-MS. The crosslinked rsNS1 alone appeared as higher oligomer bands of dimers, tetramers, and hexamers on the SDS-PAGE analysis (Fig. 4f, lane 3) which differed from the crosslinked isNS1 (Fig. 4f, lane 3). Interestingly, the addition of human HDL and crosslinker resulted in a smearing band ranging from 30-150 kDa with the loss of the well-defined monomeric rsNS1 band (Fig. 4f, lane 4). In the crosslinked rsNS1 in the absence of HDL, we identified 17 NS1:NS1 intramolecular crosslinks at the β-roll and wing domains (Fig. 4f-g, Supplementary File 3a-b). In the presence of human HDL, we identified 30 ApoA1:ApoA1, 4 NS1:NS1, and 6 NS1:ApoA1 crosslinks (Supplementary File 3a-b). Most crosslinks identified are ApoA1:ApoA1 intramolecular crosslinks in the rsNS1:ApoA1 samples (Fig. 4g). Only 1 out of 6 rsNS1:ApoA1 intermolecular crosslinks was within the 30 Å cutoff distance when mapped to the NS1:ApoA1 cryoEM model (Fig. 4e). This was in stark contrast with the isNS1 samples where the number of crosslinks between ApoA1:ApoA1, sNS1:sNS1, and sNS1:ApoA1 was similar (Fig. 4b). Overall, this suggests that rsNS1 may form oligomers of dimers but not a complex with HDL under the tested conditions.

### Ab56.2 binds at the C-terminal **β**-ladder region of NS1

The cryoEM map for isNS1wt:Fab56.2 reliably placed the epitope site of Ab/Fab56.2 at the C-terminal b-ladder region, although the precise identification of the epitope site was hindered by the limited resolution of the cryoEM maps (Fig. 2-3, Supplementary Fig. 4-7). Attempts to improve the resolution were challenging so we resorted to using a truncated form of recombinant secreted NS1 consisting of only the C-terminal region (rsNS1c; residues 172-352) to determine Ab56.2 epitope. The rsNS1c fragment of residues 172-352 was previously used for structural studies of dengue^33,34^ and other flaviviruses, West Nile virus^33^, Zika virus^34,35^, and Japanese encephalitis virus^36^. The epitope region of Ab56.2 was first confirmed by overlapping 15-mer NS1 peptide competition ELISA for Ab56.2 binding (Supplementary Fig. 9a, Supplementary Table 2). The greatest competition was observed at the NS1 peptide spanning residues 316-330 out of 301-340 which showed a decrease in OD450 absorbance (Supplementary Fig. 9b). We further applied the hydrogen-deuterium exchange mass spectrometry (HDX-MS) to monitor the conformational dynamics between Fab56.2 and NS1c to obtain higher peptide-level resolution for the binding interface. Comparative HDX results revealed global scale protection against deuterium exchange of NS1c in the presence of Fab56.2 (Fig. 5a, Supplementary Fig. 10a), indicative of reduced conformational dynamics. Large-scale differences in deuterium exchange were primarily observed across peptides spanning the dimer interface (residues 174-186) and at the tip of the C-terminal β-ladder (residues 286-347) of NS1c (Fig. 5b). Multiple overlapping peptides, spanning residues 286-347 (Fig. 5b), showed protection against deuterium exchange even at the longer labeling times (100 min) in NS1c:Fab56.2 complex (Supplementary Fig. 10a, Supplementary File 4). These results suggest that Fab56.2 bound stably with NS1c. Coupled with the observation that the most significant change in the differential plot was at residues 300-310 (Fig. 5a), these results collectively indicate that NS1c binds stably to Fab56.2 at the primary epitope site of residues 300-310.

**Fig. 5.**
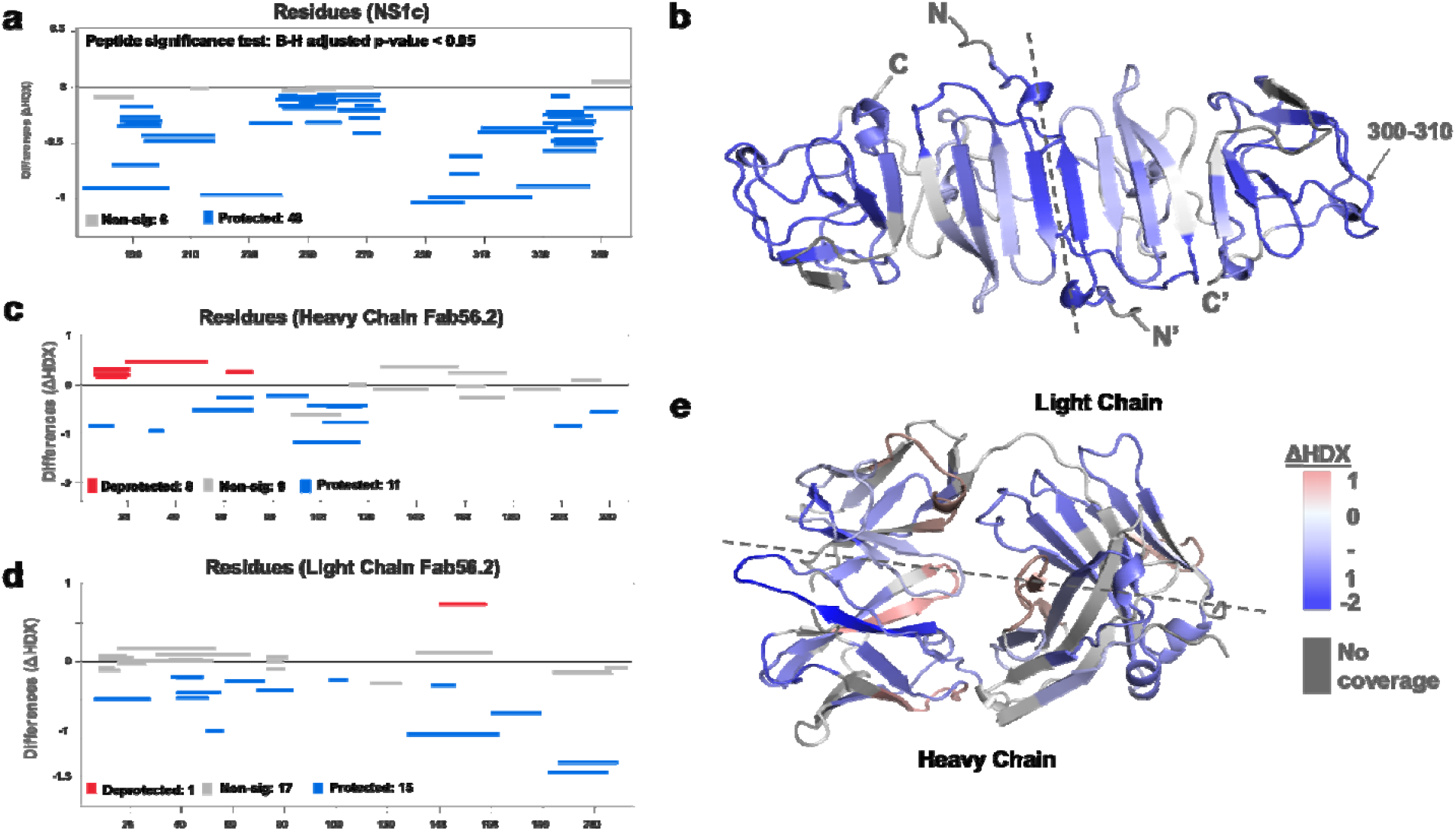
Interaction interface of rsNS1c-Fab562 complex characterized by HDX-MS. **(a)** Woods differential plot showing the differences in deuterium exchange (y-axis) between Fab562-bound and free NS1c protein across various residues (x-axis) at 10 min labeling timepoint. Negative differences indicate protection against deuterium exchange across NS1c peptides in the presence of Fab562, as compared to free NS1c. A p-value <0.05 is considered a significance threshold, which identified 6 non-significant peptides (grey lines) and 48 protected peptides (blue lines). **(b)** Cartoon representation of NS1c dimer model showing differences in deuterium exchange at 10 min labeling as indicated in key. Dashed line distinguishes the two monomers in the dimer. Peptide spanning residues 300-310 showing the highest protection are indicated. **(c, d)** Woods differential plot comparing the differences in deuterium exchange of Fab562 in the presence and absence of NS1c for various peptides of **(c)** heavy chain, and **(d)** light chain. A p-value <0.05 is considered as significance threshold, which identified deprotected (red lines), non-significant, and protected peptides as indicated. **(e)** Cartoon representation of Fab562 model showing deuterium exchange differences at 10 min mapped for Fab562-NS1c complex, as per key. Plots were generated using Deuteros 2.0, while cartoon structures were generated using PyMoL.

Comparative HDX results of NS1c-bound and free Fab56.2 revealed protection against deuterium exchange of all three complementarity-determining regions (CDR) of the heavy chain (Fig. 5c, Supplementary Fig. 10b) and CDRL2 of the light chain (Fig. 5d, Supplementary Fig. 10c). Upon binding to NS1c, peptides spanning CDRH3 (residues 89-116) showed the largest magnitude decrease in deuterium exchange at all labeling time points, while CDRH2 showed minor significant differences. In addition, only the CDRL2 loop (residues 49-56) of the light chain of Fab56.2 showed significantly decreased deuterium exchange. Mapping of the deuterium exchange profiles of heavy and light chains onto a model of Fab56.2 (Fig. 5e) revealed that CDRH1-3 and CDRL2 loops were spatially co-localized to form the paratope site to bind NS1c, with the heavy chain being the primary anchor. Collectively, these results indicated that residue 300-310 of NS1 act as the primary epitope site to bind Fab56.2 and Fab56.2 binds stably with NS1c.

### Circulating NS1 is in the form of NS1:HDL complex

Lastly, we sought to provide *in vivo* evidence of NS1 and ApoA1 interaction using sera samples from DENV-infected mice and a human patient. Since we previously showed that DENV2 T164S mutant virus led to more severe disease in the AG129 mouse model^23^, we used the same infection model to obtain pooled mice serum for immunoaffinity purification of sNS1ts as described in Supplementary Fig. 1. The mouse sera immunoaffinity purified sNS1ts was detected at ∼250 kDa by both anti-ApoA1 and anti-NS1 antibodies in a Western blot analysis following separation on a Native PAGE (Fig. 6a), supporting our mass spectrometry finding that ApoA1 is part of the ∼250 kDa isNS1 protein complex purified from infected Vero cells (Fig. 1d). We next examined if the isNS1:HDL complex can be detected in dengue patients using serum samples obtained from the CELADEN clinical trial^37^. A primary infection sample with high levels of sNS1 (>15 µg/ml) detected by commercial Platelia™ NS1 ELISA was selected for this purpose to avoid the confounding factor of the presence of anti-NS1 Abs in the secondary infected patients and to pull-down sufficient sNS1 from the patient serum. The patient serum was first subjected to a pre-clearing step using Protein AG resin to reduce the abundance of human IgGs that may interfere with the downstream analysis. We then subjected the pre-cleared patient serum to immunoprecipitation using commercial polyclonal anti-ApoA1 antibodies to pull down and detect ApoA1 and possibly sNS1. Increasing the anti-ApoA1 antibody from 10 to 50 µg coupled onto Protein AG resin for pull-down resulted in a corresponding increase in the sNS1 amount detected by ELISA (Fig. 6b), suggesting a specific association between sNS1 and ApoA1. Collectively, despite the limited amount of sNS1 in sera samples from DENV-infected mice or humans, our data points to the association of ApoA1 with sNS1 *in vivo*.

**Fig. 6.**
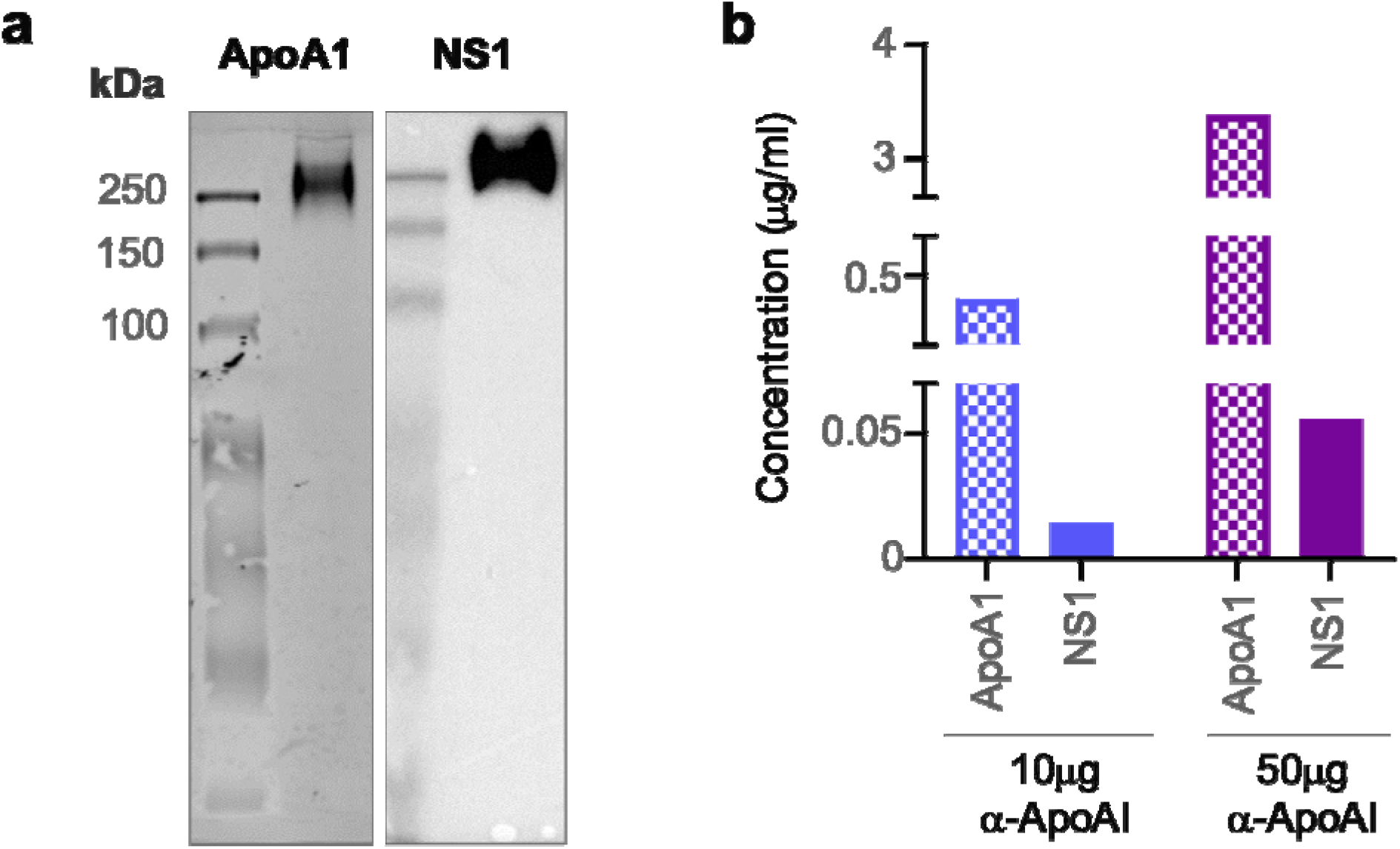
sNS1 associates with ApoA1 in DENV infected mouse and human serum. **(a)** AG129 mice (n = 10) were infected with DENV2 NS1 T164S mutant virus and the pooled infected sera collected on day 4 post-infection was subjected to NS1 immunoaffinity purification using anti-NS1 56.2 coupled resin as in Supplementary Fig. 1. 2 mg of the purified eluate was then subjected to Western blot analysis after separation and transfer from Native PAGE for detection of ApoA1 and NS1. ApoA1 was detected using the mouse monoclonal anti-ApoA1 clone 513 (Invitrogen, MIA1404) **(left panel)** and the oligomeric NS1 was detected using anti-NS1 56.2 IgG clone **(right panel)**. **(b)** Protein AG resin (Pierce) pre-cleared DENV1 infected patient serum (n = 1) from the CELADEN trial^37^ was immunoprecipitated with 10 or 50 mg of rabbit polyclonal anti-ApoA1 antibody (Biorbyt, orb10643) to detect association between ApoA1 and NS1 by ELISA. The amount of ApoA1 and NS1 in the immunoprecipitated sample was determined by human ApoA1 (Abcam, ab189576) and Platelia^TM^ NS1 Ag (Bio-Rad) ELISAs.

## Discussion

In this study, by integrating biochemical and biophysical approaches including immunoaffinity purification, cryoEM, XL-MS and HDX-MS, we have successfully purified and characterised a native population of isNS1. We determined the cryo-EM structure of a natively secreted NS1 from DENV2-infected cells as a ternary complex of NS1 dimer embedded on a single HDL particle. Our in-solution cross-linking mass spectrometry (XL-MS) analysis captured interaction within a cut-off distance of 30 Å imposed by the linker-spacer between the β-roll and wing domains of NS1 and helices 3, 5, 8, and 10 of (bovine) ApoA1 – the main protein moiety of HDL. More importantly, we were able to detect NS1:HDL complexes *in vivo* in both the sera of DENV2-infected mice and in the serum of a DENV1-infected patient, highlighting the biological relevance of the NS1:HDL complexes in dengue infection.

Benfrid et al. (2022) recently reconstituted complexes of rsNS1 dimers bound to the surface of a spheroid human HDL particle *in vitro* at 1:1 to 3:1 ratio when up to 400 μg/mL of rsNS1 were used^18^. Similarly, in our studies with infection-derived sNS1 (isNS1), we observed up to two NS1 dimers docked onto an HDL particle (Fig. 2-3, Supplementary Fig. 1c, 4-7). It is noteworthy that the concentration of isNS1:HDL in our study (∼30 μg/mL) is closer to the maximum clinically reported concentration of sNS1 in the sera of DENV patients (50 μg/mL)^3,38,39^. Therefore, the ratio of sNS1 to HDL varies depending on the concentration of the proteins used for complex formation. Furthermore, the observation that NS1 interacts with HDL regardless of the source of mammalian ApoA1 is unsurprising, given the high sequence similarity of bovine and mouse HDL to human HDL of approximately 79% and 65% respectively (Supplementary Fig. 11) and that the structure of ApoA1 is highly conserved ^40^

Acknowledging the limitation of our study that we used only one antibody (56.2) for our immunoaffinity purification, our key take-home message in terms of structure is that our data does not support the widely-held view that sNS1 is composed of a trimer of NS1 dimers with a ∼28 Å wide central channel filled with lipid as established in several seminal publications^14–16^. In fact, our data confirms the observations reported in Benfrid et al.^18^ and in the rare instances where trimer of dimers have been experimentally shown by cryoEM of rNS1 structures^17^, only a single lipid molecule can be accommodated in the narrow central channel and unlikely to be reflective of sNS1 circulating in serum. We argue that the amount of lipid associated with infection derived sNS1^15,23^ is consistent with NS1 dimer(s) associated with HDL. The interaction with HDL requires an exposed hydrophobic face of NS1 which is incompatible to the hexamer model of sNS1. Indeed, it is agreeable that the oligomeric state of NS1 is dynamic which may vary depending on the context and environment. This point requires further independent studies.

Our previous functional studies comparing isNS1wt with isNS1ts (T164S) showed that incubating immunoaffinity-purified sNS1 with human PBMCs from 3 independent human donors triggered the production of proinflammatory cytokines IL6 and TNFa in a concentration dependent manner and that it was more pronounced with the mutant than the WT^23^. Furthermore, we have provisional transendothelial electrical resistance (TEER) assay data with isNS1 proteins used in the structural studies with human umbilical vascular endothelial cells (hUVEC) which show that acid-eluted isNS1 proteins in this study induces endothelial hyperpermeability as shown in previous studies^41^. The presence of a significant population of free isNS1ts dimers in complex with Fab56.2 and fewer crosslinks identified between isNS1ts and ApoA1 point towards the possibility that isNS1ts is more dynamic in its association with host proteins than isNS1wt. The disease severity and increased complement protein expression in AG129 mice liver infected with NS1T164S carrying virus compared with WT that was observed in our previous study ^23^ can be ascribed to weakly bound mutant NS1 with fast on/off rate with HDL being transported to the liver where specific receptors bind to free sNS1 and interact with effector proteins such as complement to drive inflammation and associated pathology. This notion also requires further validation with other mutations that can impact sNS1 – HDL interaction.

Overall, our work provided a refined understanding of the native structure of sNS1 in DENV pathogenesis. The functional role of HDL-bound NS1 protein in disease is now a topic of contention but one that will be resolved by more data obtained with infection-derived sNS1 by scientists in the field. Contrary to the known protective roles of HDL such as against endotoxic shock caused by lipopolysaccharides^42–44^, Benfrid et al. (2022) showed that sNS1:HDL complex triggers proinflammatory signals when tested with primary human macrophages^18^. The presence of sNS1:HDL complexes as a predominant population of sNS1 in our *in vitro* infection model corroborates with a previous report of an interaction between insect-derived recombinant NS1 and the classical HDL/ApoA1-binding receptor known as SR-BI^19^, which is primarily expressed on hepatocytes^45^. This interaction in hepatocytes have been suggested to trigger endocytosis of sNS1 and subsequent downstream effects including vascular permeability, which is a characteristic feature of severe dengue^19,46,47^. Internalisation of sNS1 is critical for increased virus production in hepatocytes^46^ and NS1-mediated permeability in endothelial cells^47^. Furthermore, NS1:HDL complexes have been shown to induce the production of pro-inflammatory cytokines in macrophages and their presence has been detected in the sera of hospitalised DENV patients^18^. Yet, this hypothesis is confounded by other reports. While rsNS1 has been shown to trigger a TLR4-mediated release of proinflammatory cytokines to induce vascular leakage^48,49^, Coelho et al. (2021) further showed that lipid-free recombinant ApoA1 neutralizes the proinflammatory effects caused by insect-derived recombinant intracellular NS1^49^. Furthermore, exogenous administration of purified HEK-293 derived NS1 failed to worsen *in vivo* vascular leakage in sublethally infected mice^50^. This underscores the need for physiologically relevant studies to reconcile these discrepancies and gain a comprehensive understanding of the role of NS1-HDL complex in DENV pathogenesis.

In conclusion, despite the study limitations that our isNS1wt and isNS1ts were immunoaffinity purified with one specific monoclonal antibody (Ab56.2), our structural model, obtained by CryoEM and verified by XL-MS, of infection-derived sNS1 shows a NS1 dimer embedded on a single HDL particle composed of ApoA1 dimer. Future investigation of this work using infected mouse sera from other DENV strains and different flaviviruses would inform the structural and functional relevance of the HDL-bound NS1 protein complex amongst flaviviral NS1 as well as provide a basis for the development of NS1-directed therapeutics.

## Acknowledgments

We thank the scientific facility support from NTU Institute of Structural Biology and Protein Product Platform. The authors acknowledge the Cryo-Electron Microscopy Facility at Center for Bioimaging Science, Department of Biological Science, National University of Singapore the scientific and technical assistance. Recombinant DENV2 NS1 protein is a generous gift from Dr Shu Bo and Professor Lok Shee Mei. We thank members of the DL and SV labs for their support.

## Funding

This research is supported by the Singapore Ministry of Education under its Singapore Ministry of Education Academic Research Fund Tier 2 (T2EP30220-0020) to DL and National Medical Research Council of Singapore (MOH-OFIRG18may-0006; MOH-OFIRG20nov-0002) to SV, A*STAR core funding and Singapore National Research Foundation under its NRF-SIS “SingMass” scheme to R.M.S, Career Development Award 2021 (A*STAR BMRC) to W.W.P., Career Development Fund 2021 (A*STAR BMRC) to S.Z., and LKCMedicine Dean’s Postdoctoral Fellowship to C.B.L.A.

## Author contributions

Conceptualization: SV and DL; Investigation: BLAC, AQN, WWP, KWKC, ZS, SSL, MJGW, MMC, JGL, EEO, SV and DL; Writing: BLAC, AQN, SV, and DL with input from all authors.

## Competing interests

Authors declare that they have no competing interests.

## Data availability

The data that support this study are available from the corresponding authors upon reasonable request. The cryoEM density maps have been deposited in the Electron Microscopy Data Bank (EMDB) under accession codes EMD-36483 (isNS1ts:Fab56.2) and EMD-36480 (Fab56.2:isNS1ts:HDL). The protein model files for isNS1ts:Fab56.2 and Fab56.2:isNS1ts:HDL model are available upon request. Crosslinking MS raw files and the search results can be downloaded from https://repository.jpostdb.org/preview/14869768463bf85b347ac2 with the access code: 3827. The HDX-MS data is deposited to the ProteomeXchange consortium via PRIDE partner repository^51^ with the dataset identifier PXD042235.

## Notes

### Competing Interest Statement

The authors have declared no competing interest.

### Summary of Updates

Minor typo fixes.

