## Supplementary File for "Secreted dengue virus NS1 from infection is predominantly dimeric and in complex with high-density lipoprotein"

**This PDF file includes:**


Materials and Methods
Supplementary Fig. 1 to 12
Supplementary Table 1 - 2

### Materials and Methods

#### Cells, viruses, and antibodies

Vero cells (Green African monkey kidney epithelial cells, ATCC) were cultured in DMEM containing 4.5 g/L glucose (Gibco) supplemented with 10 % (v/v) fetal bovine serum (FBS) and 1 % (v/v) penicillin-streptomycin (P/S), at 37 $^{o}$C in 5 % $CO_2$. Expi293F cells were cultured in serum-free Expi293 Expression Medium at 37 $^{o}$C in 8 % $CO_2$.

The infectious clone derived DENV2 WT (GenBank accession: EU081177) and NS1:T164S mutant viruses used in this study were described previously [1].

The anti-NS1 56.2 IgG antibody (Ab56.2) was obtained from the hybridoma cell culture as described previously[2]. Briefly the Ab56.2 hybridoma cells were grown in PFHM II medium (Gibco) at 37 $^{o}$C in 5 % $CO_2$. Culture supernatants were collected every 4 days, clarified, and filtered through 0.2 μm filter membrane. Ab56.2 was purified from the supernatant through a Protein G HiTrap column (GE healthcare) using the AKTA purification system (Cytiva). The bound Ab was eluted from the Protein G column using 0.1 M glycine (pH 2.7), neutralized with 1 M Tris-HCl (pH 9.0) and dialyzed with PBS for storage at -30 $^{o}$C until use.

#### Generation and Purification of sNS1 from virus infection

Vero cells in 20 T175 flasks were infected at a multiplicity of 0.1 with DENV2 WT and T164S viruses for one hour in serum-free DMEM and subsequently replaced with 25 mL of DMEM supplemented with 2 % FBS per flask. The flasks were then incubated for 72 h at 37 $^{o}$C in 5 % $CO_2$. The crude supernatant (500 mL) was then harvested, clarified and filtered through a 0.2 μm filter membrane (Nalgene, Thermo Fisher), followed by supplementation with 0.05 % sodium azide and cOmplete EDTA-free protease inhibitor cocktail (Roche, Sigma-Aldrich). The crude supernatant was then concentrated 10-fold volume-wise using a Vivaflow 200 cassette with a 100 kDa MWCO (Sartorius) attached to a peristaltic pump (Cole Parmer). 5 mL of Aminolink™ resin (Thermo Fisher) with immobilised Ab56.2 was then added to the crude supernatant and allowed end-over-end rotation overnight at 4$^{o}$C for batch immunoaffinity purification.

The slurry was then poured into a Econo-Pac® Chromatography Column (Bio-rad), washed with filtered PBS (pH 7.4) for at least 10 column volumes and eluted with 3 column volumes of 0.1 M glycine (pH 2.7), immediately neutralised with 1 M Tris-HCl (pH 9.0). The eluted protein was dialysed against PBS (pH 7.4) overnight at 4 $^{o}$C and subsequently concentrated using a 100 kDa MWCO Amicon ultracentrifugal unit (Millipore, Merck). The total protein concentration was determined using Bradford assay (Bio-rad) and the Platelia NS1 ELISA kit (Bio-rad). Purified baculovirus-derived sNS1, as described previously[2], was used to generate a standard curve ranging from 20 ng/mL to 312.5 pg/mL for the NS1 ELISA. Protein quality was assessed on a 4-20 % polyacrylamide SDS gel and a 10 % Native polyacrylamide gel and transferred to PVDF membranes for western blot analysis against NS1 (Ab56.2) and ApoA1 (Biorbyt, orb10643). Protein purity was assessed on a 4-20 % polyacrylamide SDS gel and stained using Coomassie blue (0.2 % Coomassie blue, 7.5 % acetic acid, 50 % methanol). The Precision Plus Protein Dual Color Standard (Bio-rad) was used as the ladder for all protein gels in this work. Western blots and coomassie blue-stained gels were visualised with a Chemidoc Imager (Bio-rad). The purified protein was stored at -80 °C until use.

**Cloning and plasmid preparation of recombinant His-tagged NS1c construct**

The DENV 2 NS1c fragment was amplified from the full-length DENV2 3295 infectious clone plasmid DNA as previously described[1] using the forward primer that also includes a receptor-type tyrosine-protein phosphatase S (PTPRS) signal peptide: 5'-GGGTTGCGTAGCTGAAACCGGTAAAGAAAGGCAGGATGTATCTTGT – 3' and reverse primer: 5' - GGTGGTGCTTGGTACCGGCTGTGACCAAAGAGTTGACC – 3'. The fragment was then cloned into the 6xHis sequence-containing pHL-sec vector at XbaI/KpnI cloning sites. The plasmid was then amplified in XL1-blue *Escherichia coli* and plasmid preparation was performed using the Plasmid DNA Maxiprep kit (Thermo Fisher).

**Generation and purification of recombinant His-tagged NS1c**

Expi293F cells were transfected at a cell density of 3 x $10^6$ viable cells/mL and with 1 µg of the NS1c plasmid DNA/mL culture volume, using the ExpiFectamine 293 Transfection kit (Gibco, Thermo Fisher), according to manufacturer's instructions. On day 5 post transfection, the cells were spun down and the supernatant containing the expressed His-tagged NS1c was harvested, clarified, and filtered using a 0.22 um filter membrane (Millipore, Merck). The His-tagged NS1c was then purified from the supernatant using a 1 mL HisTrap column (Cytivia), concentrated using a 50 kDa MWCO Amicon concentrator (Millipore, Merck) and stored in –80 $^oC$ until use. Protein yield and quality was assessed on a 4-20 % reducing SDS-PAGE and stained with Coomassie blue, as described in **Generation and Purification of sNS1 from virus infection**.

**Generation and purification of anti-NS1 56.2 Fab**

1.8 mg of the purified Ab56.2 was subjected to papain digestion using the immobilized papain resin (Thermo Fisher) at an enzyme:substrate ratio of 1:160, according to the manufacturer's instructions. The IgG-papain mixture was incubated at 37 $^oC$ for 3.5 hours followed by removal of the immobilized papain resin to stop the digestion. The papain IgG digest was buffer-exchanged to PBS using PD10 Sephadex G25 column (GE healthcare). The resulting Fab (Fab56.2) was then purified from the crude digest by Protein A HiTrap column (GE healthcare) using the AKTA purification system (Cytiva), concentrated using 3 kDa MWCO Amicon concentrator (Millipore, Merck) and stored at -30 $^oC$ until use.

**Analytical size exclusion chromatography for isNS1, Fab and isNS1:Fab complexes**

10 µg of isNS1wt or isNS1ts was complexed with papain-digested Fab56.2 or Ab56.2 at a protein:Fab or protein:Ab molar ratio of 1:5 by incubation for at least 2 hours on ice. 10 µg of isNS1wt, isNS1ts or papain-digested Fab56.2 as well as the isNS1wt: or isNS1ts:Fab56.2 complex was then independently injected onto a Superdex 200 increase 3.2/300 GL column (GE Healthcare) connected to the AKTA purification system (Cytiva) in PBS (pH 7.4) at a constant flow rate of 0.075 mL/min. Chromatograms were analyzed on Unicorn 7 and replotted on OriginPro, Version 2021b.

**Negative stain microscopy and data processing**

3 µL of protein sample at concentration of 0.01 mg/mL were spotted on glow-discharged carbon grids, contrasted with 2 % uranyl acetate, and imaged with a FEI Tecnai T12 microscope equipped with an Eagle 4 megapixel CCD camera (Thermo Fisher, USA). 24 micrographs were manually collected and processed using Scipion3[3]. CTF Estimation,

manual and auto-picking were done using xmipp3[4] resulting in a total of 3914 particles. This is followed by particle extraction and 2D classification using Relion[5].

**Cryo-EM grid preparation and microscopy**

QuantiFoil or UltrAuFoil R1.2/1.3 gold 300 mesh grid was covered with a graphene layer (Graphenea) following an adapted protocol[6] and glow-discharged for 10 seconds at low energy (Harrick Basic Plasma Cleaner) before use. 2.5 µL of protein sample at concentration of 0.35 mg/mL was applied to the grids, blotted for 3 seconds with blot force 1, and plunge-frozen in liquid ethane using Vitrobot (Thermo Fisher Scientific). Cryo-EM grids for isNS1, isNS1wt:Ab562, and isNS1wt:Fab562 were imaged using EPU v2.14 on a 300 kV TEM Titan Krios with Fringe-free imaging and aberration-free image shift (Thermo Fisher Scientific) and equipped with a K2 direct electron detector (Gatan). Data collection was performed at a nominal magnification of 165,000x with a physical pixel size of 0.85 Å. A GIF Quantum Energy Filter with a slit width of 20 eV was used. Micrographs were dose-fractioned into 40-50 frames with a total exposure dose of 53-70 e- per Å^2. The isNS1ts mutant in complex with Fab56.2 dataset was collected on a Titan Krios equipped with a K3 direct electron detector (Gatan) using SerialEM v4.06 [7]. Data collection was performed at a nominal magnification of 105,000x with a physical pixel size of 0.858 Å. A GIF Quantum Energy Filter with a slit width of 20 eV was used. Micrographs were dose-fractioned into 40 frames with a total exposure dose of 53 e- per Å^2. The data collection parameters are summarized in Supplementary Table 1.

**Cryo-EM image processing and model fitting**

Collected movies were imported to Cryosparc v3.3 or later[8] and processed similarly for all datasets starting with Patch-Based Motion Correction and CTF Estimation. Micrographs with CTF-estimated maximum resolution better than 4.5 Å were selected for particle picking starting with blob-based picker followed by a combination of Template Picker, Topaz[9] and crYOLO v1.7.6[10] after initial rounds of 2D classification using extracted particles that were fourier cropped by four times had been performed. Particles were extracted at a box size at 300 pixels for isNS1ts (Supplementary Figure 4), 352 pixels isNS1wt:Ab562 (Supplementary Figure 5), and 416 pixels for isNS1wt:Fab562 (Supplementary Figure 6) and isNS1ts:Fab562 (Supplementary Figure 7). Additional two rounds of 2D classification without downsampling was performed before the selected particles were subjected to ab initio reconstruction for 3 classes. All resulting classes were refined respectively with no symmetry (C1) using Heterogeneous Refinement for further cleaning followed by Non-Uniform Refinement[11]. In general, all the datasets led to low resolution maps and the class maps of isNS1wt:Ab562, isNS1wt:Fab562, and isNS1ts:Fab562 could be fitted with the NS1, Fab56.2, and apoA1 models generated using Alphafold2[12,13] (Figure 2d, 3d-e). The final map used for isNS1wt:Ab562 (Supplementary Figure 5) for the overall shape and size is derived from Heterogeneous Refinement as the Non-Uniform Refinement resulted in badly connected maps (Supplementary Figure 4b). Preferred orientation is most apparent for the class maps of isNS1ts:Fab562. Further Local Refinement was attempted for selected classes of isNS1wt:Fab562 (Supplementary Figure 6) and isNS1ts:Fab562 (Supplementary Figure 7) datasets to improve final maps, however there was no visible improvement. Histogram and directional FSC plots were generated for selected classes of isNS1wt:Ab562 (Supplementary Figure 5), isNS1wt:Fab562

(Supplementary Figure 6) and isNS1ts:Fab562 (Supplementary Figure 7) using the remote 3DFSC processing server[14] for better estimation of the map resolution and directional anisotropy. Map fitting and figures representing the map and model features were performed using UCSF ChimeraX[15]. The data collection statistics are summarized in Supplementary Table 1. Data processing is also outlined in the Supplementary Figures 4 – 7.

**In gel protein identification and quantification by liquid chromatography mass spectrometry (LC-MS)**

5 µg of isNS1wt and isNS1ts were separated on a 12 % polyacrylamide SDS gel and a 10 % Native polyacrylamide gel and subjected to gel electrophoresis. The gel was then rinsed with MilliQ water before staining with Coomassie blue stain (0.2 % Coomassie blue R-250, 7.5 % acetic acid, 50 % Methanol) for 30 minutes with gentle agitation at room temperature. The gel was then destained with Coomassie blue destaining solution (40 % Methanol, 10 % acetic acid) and rinsed with MilliQ water before visualising on a Chemidoc Imager (Bio-rad).

The destained gel bands were excised and cut into smaller pieces (approximate 1 mm x 1 mm size). The gel pieces were washed with 1 ml of wash buffer 1 (50 % ethanol, 50 mM Triethylbicarbonate (TEAB) pH 8.5) (Sigma) by gentle agitation at room temperature for 5 minutes. The process was repeated twice. The gel pieces were washed twice with 1 ml of wash buffer 2 (100 % ethanol) for 5 minutes. The ethanol was removed, and the gel pieces were resuspended in 100µL 100 mM TEAB pH 8.5. The samples were subsequently reduced with 10mM Tris(2-carboxyethyl)phosphine hydrochloride (TCEP) (Sigma) by incubating with 20 minutes at 25 °C shaking and were alkylated with 55mM Chloroacetamide (CAA) in the dark for 20 minutes at 25 °C shaking. The proteins were digested by incubating with 0.75 µg of trypsin (Thermo Fisher Scientific) at 25 °C overnight. The supernatant was removed, and the digested peptides were extracted using 100 µL of 30 % acetonitrile (ACN) with 3 % formic acid (FA) and 100µL of 80 % acetonitrile (ACN) with 3 % formic acid (FA). Each extraction step was repeated twice. The digested peptides were dried and resuspended in 20 µL A*buffer (2 % ACN, 0.06 % Trifluoroacetic acid, 0.5 % acetic acid) and 2 µL of samples were injected to LC-MS for protein identification and quantification. The peptides were separated on EasySprayTM (Thermo Fisher Scientific) column with 75 µm inner diameter 50 mm length with particle size 2 µm and a pore size of 100 Å over a 45-minute gradient starting with buffer A containing 0.5 % (v/v) formic acid to buffer B containing 95 % acetonitrile with 0.5 % (v/v) formic acid with nanoflow LC pump Easy nLC1200 with a flow rate of 300 nL per minute. The data was acquired on Thermo Fisher Orbitrap HF-X mass analyzer or Thermo Fisher Orbitrap Lumos mass analyzer in data dependent acquisition mode with each MS scan followed by MS/MS scans. Each duty cycle is 2.5 seconds long with MS scan at 60,000 m/z resolution followed by MS/MS scans at 7,500 m/z resolution with AGC target of $4x10^4$ and dynamic exclusion of 30 seconds. Raw files were analysed and quantified using proteome discoverer (PD), version 2.4 (Thermo Fisher Scientific). The proteins were identified using MASCOT search engine using UniProt Bovine Proteome and NS1 protein sequence from DENV2 Singapore clinical isolate with Genbank accession number EU081177.1 with following parameters, trypsin as the digesting enzyme with maximum two mis-cleavages. The oxidation of methionines, acetylation of protein N-termini, deamidation, were set as variable modifications and carbamidomethylation of cysteines were set as the static modification. The MS tolerance was set at 15 ppm, MS/MS tolerance at 0.08 Da. The precursor ions were quantified using both unique + razor peptides.

**Cross-linking Mass Spectrometry**

Purified isNS1wt complex and isNS1ts were independently crosslinked using disuccinimidyl sulfoxide (DSSO) (Thermo Scientific) in PBS, according to the manufacturer's instructions. Briefly, 2.4 µM (or 0.6 mg/mL) of total protein in the purified isNS1wt and 5.8 µM (or 1.45 mg/mL) of total protein in the purified isNS1ts were crosslinked with DSSO in an isNS1:DSSO molar ratio of 1:100 for 45 minutes at room temperature. The crosslinking reaction was then quenched with 20 mM Tris-HCl (pH 8.0) for 15 minutes at room temperature. The crosslinked proteins were assessed on a 4-20 % reducing SDS-PAGE before subjecting to a silver stain, according to manufacturer's instructions (Pierce) and a Western blot using the anti-NS1 antibody Ab56.2.

Commercial DENV2 sNS1 (2.7 µM or 0.675 mg/mL) (Native Antigen Company) and commercial purified HDL (7.1 mM or 2.3 mg/mL) (Innovative Research) in PBS were allowed to associate by incubation on ice for 2 hours in a molar ratio of 1:1. DSSO crosslinker was then added in 100-fold molar excess. Crosslinking was allowed to proceed for 45 minutes at room temperature before quenching with 20 mM Tris-HCl (pH 8.0) for 15 minutes at room temperature. The crosslinked products were assessed on a 4-20 % reducing SDS-PAGE before subjecting to a silver stain, according to manufacturer's instructions (Pierce) and a Western blot using the anti-NS1 antibody Ab56.2.

The crosslinked proteins were resuspended in 45 µL of 100 mM TEAB, reduced and alkylated with 10mM of TCEP and 55 mM of CAA. The proteins were digested by incubation with trypsin (Thermo Fisher Scientific) overnight at 25 °C. The digested peptides were acidified and desalted on stage tips packed with 3M Empore C18 disc. The peptides were eluted with elution buffer (80 % ACN 0.5 % acetic acid) and dried. The desalted peptides were resuspended in 10 µL of A*buffer. 1 µg of digested peptides were separated on an EasySpray (Thermo Fisher Scientific) column with 75 µm inner diameter 50 mm length with particle size 2 µm and a pore size of 100 Å over a 75-minute gradient starting with buffer A containing 0.5 % (v/v) formic acid to buffer B containing 95 % acetonitrile with 0.5 % (v/v) formic acid with nanoflow LC pump Easy nLC1200 with a flow rate of 300 nL per minute. The data was acquired on an Orbitrap Fusion Lumos mass spectrometer (Thermo Fischer Scientific) with CID-MS2/HCD-MS3 fragmentation. MS1 scans were performed in the Orbitrap with 60,000 m/z resolution, AGC target of 400,000 and maximum injection time of 35 ms. Precursor ions with positive charge state 3-8 were selected for MS2 fragmentation with dynamic exclusion of 60 s after 1 scan. MS2 scans were acquired in the Orbitrap with 30,000 m/z resolution, AGC target of 50,000, maximum injection time of 50ms and CID collision energy of 30 %. Targeted mass difference of 31.9721 m/z (signature peak of MS-cleaved DSSO) were selected for MS3 acquisition in the Ion Trap with rapid scan rate, AGC target of 10,000, maximum injection time of 40 ms and HCD collision energy of 30 %. Raw files were searched using Metamorpheus ver 0.0.320 [16], against a fasta database containing the amino acid sequences of DENV2 NS1

,and bovine ApoA1 for isNS1wt and isNS1ts sapmles and human ApoA1 for rsNS1 samples, with a calibration task before searching for cross-linked peptides. Calibrate task was performed with precursor mass tolerance of 10 ppm, and product mass tolerance of 20 ppm. Cross-link search was performed for DSSO cross-links at K,S,T,Y amino acid sites, with CID MS2 dissociation type and HCD MS3 child scan dissociation. Up to 3 missed cleavages were allowed and protease was set to trypsin, with precursor mass tolerance of 10 ppm and product mass tolerance of 20 ppm. Fixed modifications were set for carbamidomethylation (C), while variable modifications were set for oxidation (M), deamidation (N,Q) and for DSSO (K,S,T,Y, protein N-terminus). Resultant cross-linked peptides were filtered for q-value ≤ 0.01 (corresponds to 1% false discovery rate). Cross-link sequence representation figures were made using xiVIEW[17] while cross-links were mapped to structures using PyXlinkViewer[18] in Pymol (Schrödinger, LLC.) and XL Mapping and Analysis (XMAS)[19] with UCSF ChimeraX [15].

**Immunoprecipitation of ApoA1-sNS1 complex from DENV infected patient serum**

To demonstrate the physiological relevance of the sNS1:ApoA1 interaction *in vivo*, a co-immunoprecipitation of sNS1:ApoA1 was performed on a DENV1 primary infected patient serum obtained from the Celgosivir clinical trial conducted in Singapore (CIRB Ref:2012/025/E) that contains ~30 μg/mL sNS1 quantified by Platelia NS1 Capture ELISA (Bio-Rad)[20]. 75 µl of AG agarose beads was pre-coupled with either 10µg or 50µg of rabbit polyclonal ApoA1 Ab (Biorbyt, orb10643). Prior to immunoprecipitation, the patient serum sample was diluted 2× with PBS and subjected to three rounds of Protein AG agarose beads (Pierce) pre-clearing to remove the highly abundant IgGs in the serum. The Ab pre-coupled AG beads were then added to pre-cleared serum sample and incubated overnight with end-to-end rotation at 4 °C. The sample-bound AG beads were washed four times with PBS-T (0.1 % Tween 20 (v/v)) followed by addition of 60 µl of 0.1 M glycine (pH 2.7) to elute the sample from the resin that was subsequently neutralized with 1 M Tris-HCl (pH 9.0). The ApoA1 and sNS1 in the eluted sample was subjected to quantification by ELISAs (commercial human ApoA1 ELISA, Abcam; Platelia NS1 Capture ELISA, Bio-Rad). A human naïve serum (obtained accordance with National University of Singapore IRB approval B-12-227.) that served as a negative control was processed and analyzed similarly as the patient serum.

**Immunoaffinity purification of sNS1 from infected mouse serum**

Sv/129 mice deficient of Type I and II IFN receptor (AG129) purchased from B&K Universal (UK), were housed in BSL-2 animal facility in Duke-NUS, Singapore. All animal experiments (protocol 2021/SHS/1646) were approved by the Institutional Animal Care And Use Committee at Singapore Health Services and conformed to the National Institutes of Health (NIH) guidelines and public law. 10 AG129 (12-14 week old male) mice were pre-injected with 50μg 4G2 antibody intravenously (i.v.) prior to intravenous inoculation of $2 \times 10^7$ pfu of DENV2 NS1 T164S mutant virus[1]. The mice were sacrificed on day 4 post-infection by $CO_2$ inhalation for blood collection from the postcaval vein. A total of 6ml serum was collected, concentrated and buffer-exchanged with PBS using 100kDa cutoff Amicon concentrator (Millipore). Finally, ~5ml of the buffer-exchanged serum was incubated with 1ml of the anti-NS1 56.2 coupled resin for immunoaffinity purification as described (see **Generation and Purification of sNS1 from virus infection** above). The elutes were concentrated using a 30kDa Amicon concentrator and subjected to ELISA quantification using commercial Platelia™ Dengue NS1 Ag kit (Bio-Rad) and Western blot analysis after separation by native PAGE.

**NS1 peptide competition ELISA**

The epitope of anti-NS1 Ab56.2 was determined using peptide competition ELISA as previously described[21]. A schematic on the principle of NS1 peptide competition ELISA is shown in Supplementary Figure 11a. An array of overlapping 15-mer peptides spanning DENV2 NS1 C-terminal residues 241-352 (GenBank accession: EU081177) were synthesized and purchased from GL Biochem (Shanghai) Ltd. 2nM of Ab56.2 was preincubated with 5 μM (2500× molar excess) for 30 minutes at room temperature before transferring to an immunoplate pre-coated with DENV2 full-length NS1 (5μg/mL in 0.1M $NaHCO_3$ buffer at pH 9.6). The immunoplate with the peptide-Ab mix was incubated at room temperature for 10 minutes followed by washing with PBS-T (0.1% v/v Tween 20). For assay controls, a 15-mer NS1 peptide spanning the β-roll residues 6-20 was used as a non-competing peptide while the full-length DENV2 NS1 was used as a competing reagent. Bound Ab was detected with an anti-mouse HRP and absorbance readings at 450 nm were measured in duplicates. Results were tabulated as mean percentage absorbance reading normalized to the Ab only control (no peptides).

**Hydrogen-Deuterium Exchange Mass Spectrometry (HDXMS)**

HDXMS was used to map the epitope and paratope sites of NS1c and Fab562 complexation, respectively. HDX was performed for NS1c and Fab562 individual purified proteins, and NS1c-Fab562 complex obtained from size-exclusion chromatography elution. To initiate the hydrogen-deuterium exchange reaction, ~75 pmol of the protein samples was diluted with phosphate-buffered saline prepared in deuterium oxide (D2O, Cambridge Isotopes) to achieve a 90% final deuteration. The deuterium exchange was carried out by incubating the proteins for 1, 10, 100 min labeling timepoints at 25 °C. After the desired labeling time, the reaction was stopped by lowering the pH to 2.5, temperature to 0 °C using a chilled quench solution (1 M Guanidinium Hydrochloride, 0.1 M TCEP). For reference, non-deuterated controls were also carried out by diluting the protein samples in aqueous PBS, followed by quench solution and mass spectrometry analysis.

Each quenched sample was subjected to 3 min proteolytic digestion using an immobilized pepsin cartridge (Enzymate$^{TM}$, Waters, USA) maintained at 12 °C, and built into an HDX sample manager coupled to nanoACQUITY M-class UPLC (Waters, USA). The samples were pumped by 0.1 % formic acid solution at 100 μl/min. The pepsin-digested peptides were then resolved by reverse-phased liquid chromatography using a C18 trap (Vanguard, Waters, USA) and a C18 column (ACQUITYTM, Waters, USA) maintained at 0-3 °C [22]. A gradient of 8-40% solvent A (0.1% formic acid in LCMS-grade water) to solvent B (0.1% formic acid in Acetonitrile) pumped at 40 μl/min by M-class binary solvent manager (Waters, USA) was used to separate the peptides and identified through a coupled high-resolution Synapt G2-Si (Waters, UK) mass spectrometer. The peptides are ionized by electrospray ionization and detected in positive polarity mode. Ionized peptides were separated using HDMS$^E$ mode with ion-mobility activated with the following parameters: low collision energy – trap (2 V), transfer (4 V); high collision energy – trap (4 V), transfer ramp (15-40 V); IMS wave velocity – 600 m/s, transfer wave velocity – 197 m/s; step-wave height at 30 V. A capillary voltage of 3 kV and 80 °C source temperature were applied for ionization.  to detect the peptides and measure their absolute masses. For mass accuracy, 10 µl/min of 200 fmol/µl of Glu-fibrinogen peptide B was simultaneously sprayed as lockmass reference.

The mass spectrometry raw data acquired was processed using Protein Lynx Global Server v3.0 (Waters, USA) to identify and annotate the peptides. Non-deuterated samples were used to match against the respective amino acid sequences of NS1c and Fab562 as the search database. The following search parameters were used for accurate matching and annotation: non-specific protease digestion, number of matches within peptide – 3, within protein – 7, a false positive rate of 4%, and a mass error of 1 ppm for precursor peptide. The annotated peptide list obtained was then filtered and analyzed using DynamX 3.0 (Waters, USA) software. Peptides with a minimum intensity of 1000, minimum products per amino acid 0.1, and a $MH^+$ error of 10 ppm were used to filter out the poor quality and low scoring peptides. This peptide list obtained from the non-deuterated controls was then used for deuterium exchange analysis using DynamX 3.0. The deuterium exchange data analyzed for the peptides in different sample conditions and across all time points was manually verified. Deuterium uptake was calculated as the difference in the masses of the centroids of the deuterated state and the corresponding non-deuterated control. The differences in deuterium exchange values between two states of the proteins were then checked for statistical significance using Deuteros 2.0 [23]. All peptides with p-value <0.05 were only considered significant. The final analysis yielded 54 peptides covering 95% of NS1c, 31 peptides covering 94% of heavy chain of Fab562, and 33 peptides covering 92.5% of the light chain of Fab562. All HDX experiments were carried out in triplicate independent measurements, and the average deuterium exchange data is tabulated in Supplementary information. Raw files are accessible via this link https://repository.jpostdb.org/preview/14869768463bf85b347ac2 with the access code: 3827.

### Supplementary figures

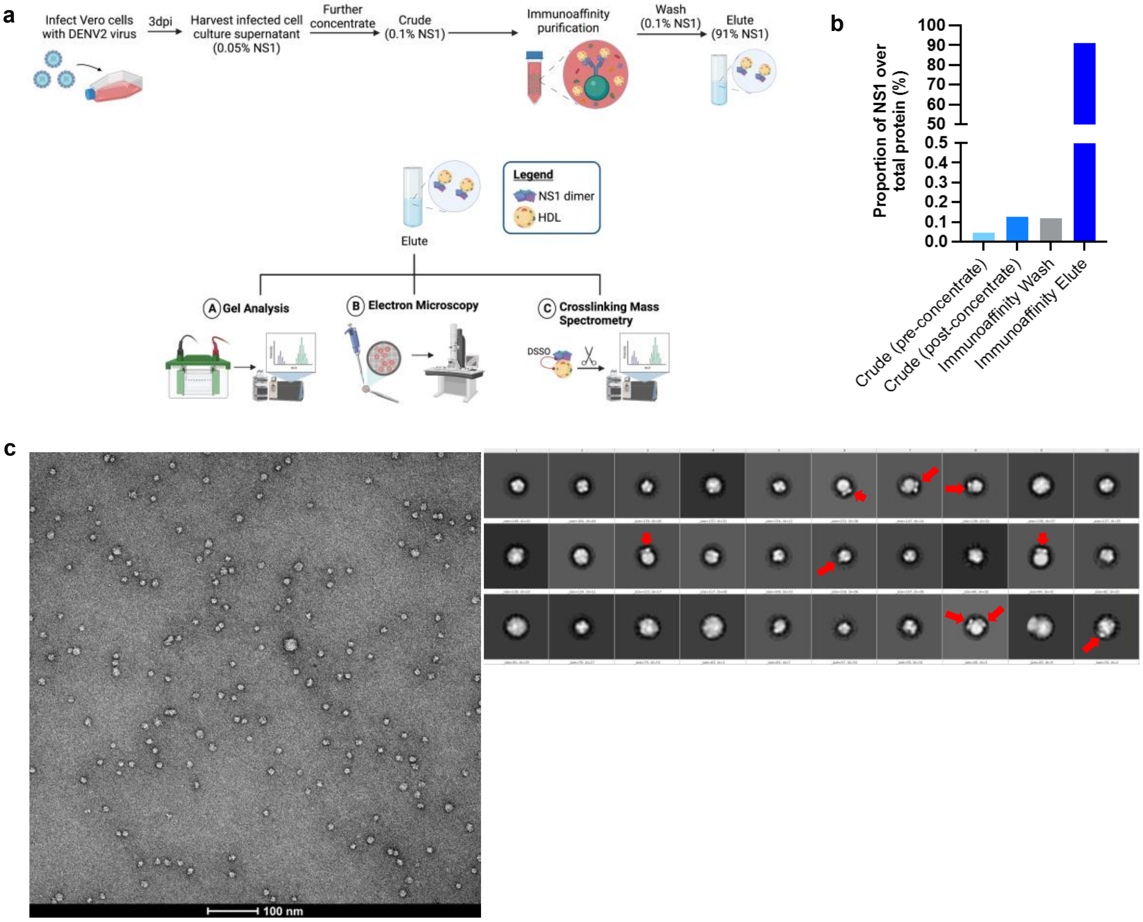


**Supplementary Fig. 1. Purification and negative stain electron microscopy screening of in vitro infection-derived sNS1 from infected Vero cells. (a)** Schematic of the isNS1 batch immunoaffinity purification protocol and the downstream analyses used in this paper (created using Biorender.com). Infected cell supernatant from Vero cells (either WT or T164S EDEN2) was harvested at 72 hpi, clarified, filtered, supplemented with a protease inhibitor cocktail and 0.05% sodium azide and finally concentrated using a 100 kDa MWCO Vivaflow casette attached to a peristaltic pump. isNS1wt or isNS1ts was then batch immunoaffinity purified using the 56.2 anti-NS1 antibody immobilised on the AminoLink™ resin. The resin was then loaded into a column, washed with at least 10 CV of PBS (pH 7.4), eluted with 0.1M glycine (pH 2.7) and immediately neutralised with 1M Tris-HCl (pH 9.0). The eluted protein was then dialysed against PBS and concentrated using a 100kDa MWCO Amicon ultracentrifugal unit and stored at –80°C before use. The protein purity was determined via Coomassie blue after separation on a reducing SDS-PAGE. The protein bands observed on the gel were then validated in a western blot against NS1 and ApoA1. Protein quality was also determined via NS1 western blot following separation on a Native-PAGE. Excised bands corresponding to 250 kDa on the Native gel, and 50 kDa and 25 kDa bands on the denatured gel as well as the Elute in solution were also subjected to protein identification via LC-MS. Purified isNS1wt and isNS1ts were separately complexed with Fab56.2 and Ab56.2 and analysed through an analytical size exclusion chromatography to ensure formation of stable complexes for imaging via electron microscopy. Purified isNS1wt and isNS1ts were also crosslinked with DSSO to determine interaction sites between isNS1 and ApoA1 via LC-MS. **(b)** Enrichment of NS1 during the immunoaffinity purification process. sNS1wt purification is used as a representative for the sNS1 purification process. The proportion of NS1 over total protein was measured by taking a percentage of the total amount of NS1 measured (using NS1 ELISA) out of the total protein measured (using Bradford assay) in the Crude supernatant before concentrating, Crude supernatant after concentrating, immunoaffinity PBS wash and finally in the immunoaffinity Elute after buffer exchange against PBS and further concentrating. There is an approximately 455-fold enrichment of NS1 from the concentrated crude supernatant to the immunoaffinity Elute. **(c)** Representative negative stain electron micrograph of isNS1 image on a 120 kV FEI Tecnai T12 equipped with an Eagle 4 mega pixel CCD camera. The corresponding 2D classes from the particles picked are shown on the right, red arrows highlighting the NS1 dimer protruding out of the spherical density.

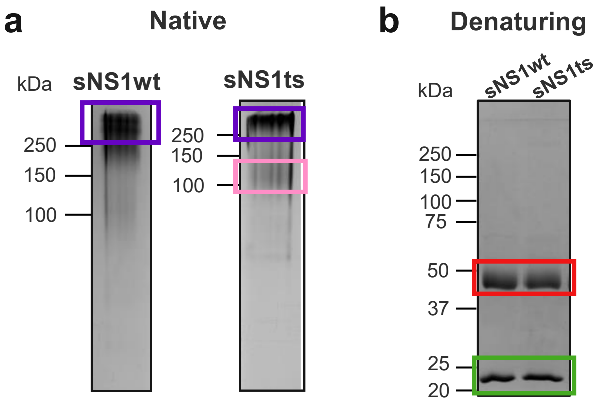


**c**

| sNS1wt | Normalized LFQ Intensities ($10^{10}$) | | | |
|---|---|---|---|---|
| | Elute | 250 kDa **Native** gel band | 50 kDa **Denaturing** gel band | 25 kDa **Denaturing** gel band |
| **NS1** | 8.73 ± 0.16 | 8.39 ± 0.09 | 166 ± 0.02 | 0.13 ± 0.04 |
| **ApoA1** | 4.65 ± 0.02 | 6.95 ± 0.06 | 0.13 ± 0.009 | 16.5 ± 0.039 |
| **Median LFQ intensity** | 0.007 ± 0.002 | 0.008 ± 0.0001 | 0.002 ± 0.0001 | 0.0007 ± 0.0001 |
| **% sNS1 intensity in TIC** | 50.61 ± 1.04 | 48.46 ± 0.64 | 96.85 ± 0.02 | 0.77 ± 0.2 |
| **% ApoA1 intensity in TIC** | 26.97 ± 1.24 | 40.1 ± 0.47 | 0.78 ± 0.05 | 96.23 ± 0.27 |

| sNS1ts | Normalized LFQ Intensities ($10^{9}$) | | | | |
|---|---|---|---|---|---|
| | Elute | 250 kDa **Native** gel band | 100 kDa **Native** gel band | 50 kDa **Denaturing** gel band | 25 kDa **Denaturing** gel band |
| **NS1** | 35.3 ± 0.24 | 63.8 ± 5.93 | 2.52 ± 0.97 | 138 ± 0.43 | 1.36 ± 0.57 |
| **ApoA1** | 50.5 ± 3.18 | 66.3 ± 7.50 | 119 ± 0.71 | 0.82 ± 0.24 | 137 ± 43.10 |
| **Median LFQ intensity** | 0.069 ± 0.039 | 0.019 ± 0.008 | 0.006 ± 0.001 | 0.003 ± 0.001 | 0.01 ± 0.003 |
| **% NS1 intensity in TIC** | 24.63 ± 0.17 | 44.51 ± 4.14 | 2.24 ± 1.76 | 96.50 ± 0.30 | 0.95 ± 0.40 |
| **% ApoA1 intensity in TIC** | 35.19 ± 2.22 | 46.25 ± 5.23 | 82.90 ± 0.5 | 0.57 ± 0.17 | 96.67 ± 0.36 |

**Supplementary Fig. 2.** Protein identification analysis of excised gel bands via LC-MS. **(a)** Coomassie blue detection of proteins of immunoaffinity-purified isNS1wt and isNS1ts (Elute) after separation on a 10% Native-PAGE. The 250 kDa bands (purple) that were present in both isNS1wt and isNS1ts, as well as the 100 kDa band (pink) that was present only in isNS1ts, were excised for protein identification analyses via LC-MS. **(b)** Coomassie blue detection of proteins of immunoaffinity-purified isNS1wt and isNS1ts (Elute) after separation on a 10% reducing SDS-PAGE. The 50 kDa bands (red) and the 25 kDa bands (green) in both isNS1wt and isNS1ts Elutes were excised for protein identification analyses via LC-MS. **(c)** Label-free quantification (LFQ) of NS1, ApoA1 and other unidentified proteins in total ion intensity via LC-MS (n=3) in the following samples fromi sNS1wt (top table) and isNS1ts (bottom table): Elute in solution, 250 kDa gel band (boxed in purple), 100 kDa gel band (boxed in pink; present only in isNS1ts), 50 kDa gel band (boxed in red) and 25 kDa gel band (green).

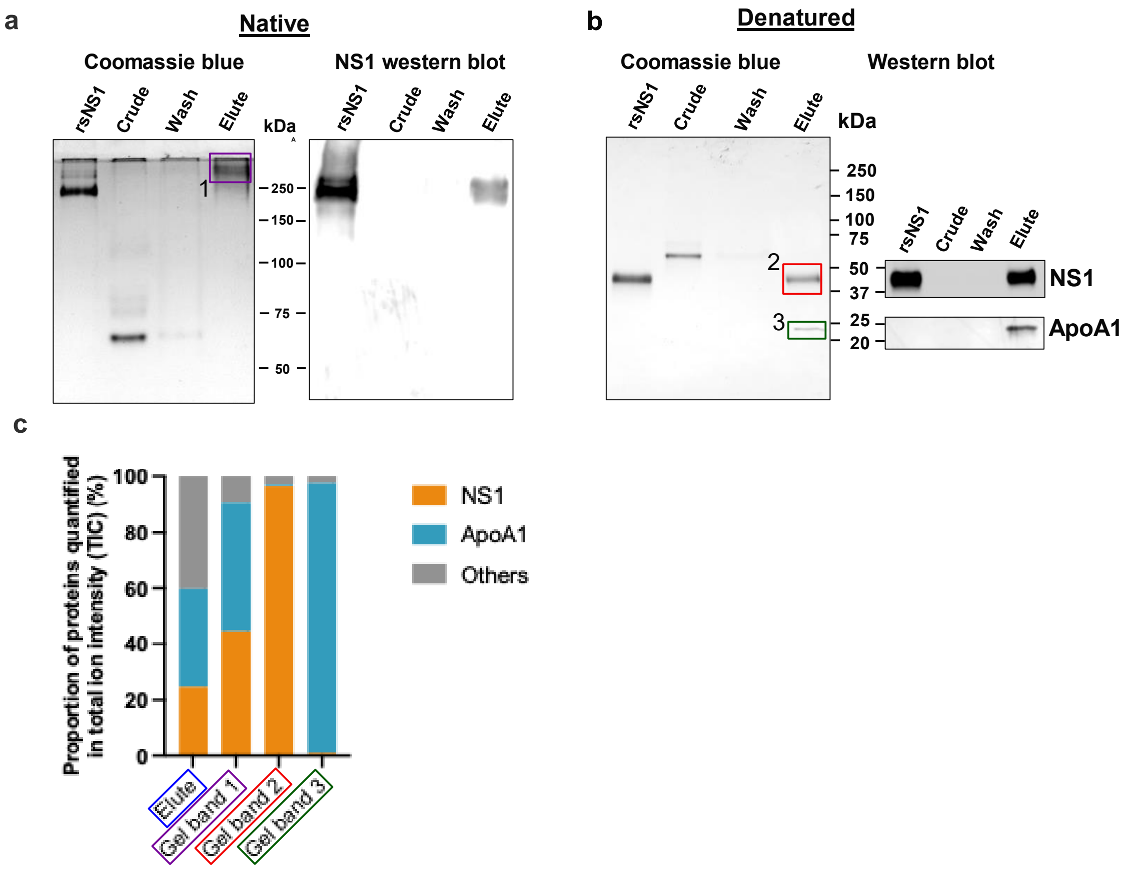


**Supplementary Fig. 3. Composition of the secreted isNS1ts from DENV-infected Vero cells. (a)** Coomassie blue detection of proteins from 1 µg of total protein of Crude and Elute, and 100 ng of total protein (in maximum well volume) in Wash immunoaffinity fractions (low concentration due to large volume), with the recombinant sNS1 (rsNS1) obtained from Shu et al (2022) as a positive control, after separation on a 10 % Native-PAGE gel (left). The same set of samples were also subjected to a western blot detection of NS1 using Ab56.2, in 500 ng of total protein (except for Wash where 100 ng in the maximum well volume) after separation on a 10 % Native-PAGE (right). **(b)** Coomassie blue detection of proteins from 1 µg of total protein of Crude and Elute, and 100 ng of total protein (maximum well volume) in Wash immunoaffinity fractions for isNS1ts and rsNS1 (Shu et al., 2022), after separation on a 4-20 % reducing SDS-PAGE gel. 1 µg of total protein in the same set of samples (except for Wash where 100ng in the maximum well volume) were also subjected to a western blot detection of NS1 and ApoA1 using Ab56.2 or ApoA1 antibody (Biorbyt, orb10643) respectively, after separation on a 4-20 % reducing SDS-PAGE (right). **(c)** In-gel protein identification of the purified isNS1ts by liquid chromatography mass spectrometry (LC-MS). Proportion of NS1, ApoA1 and other unidentified proteins quantified in total ion intensity, obtained from the following samples: Elute in solution (boxed in blue), 250 kDa gel band (boxed in purple), 50 kDa gel band (boxed in red) and 25 kDa gel band (green). The boxed gel bands are from representative gels showing the different protein species found while the actual gel bands used for protein identification by LC-MS are as shown in Supplementary Fig. S2.

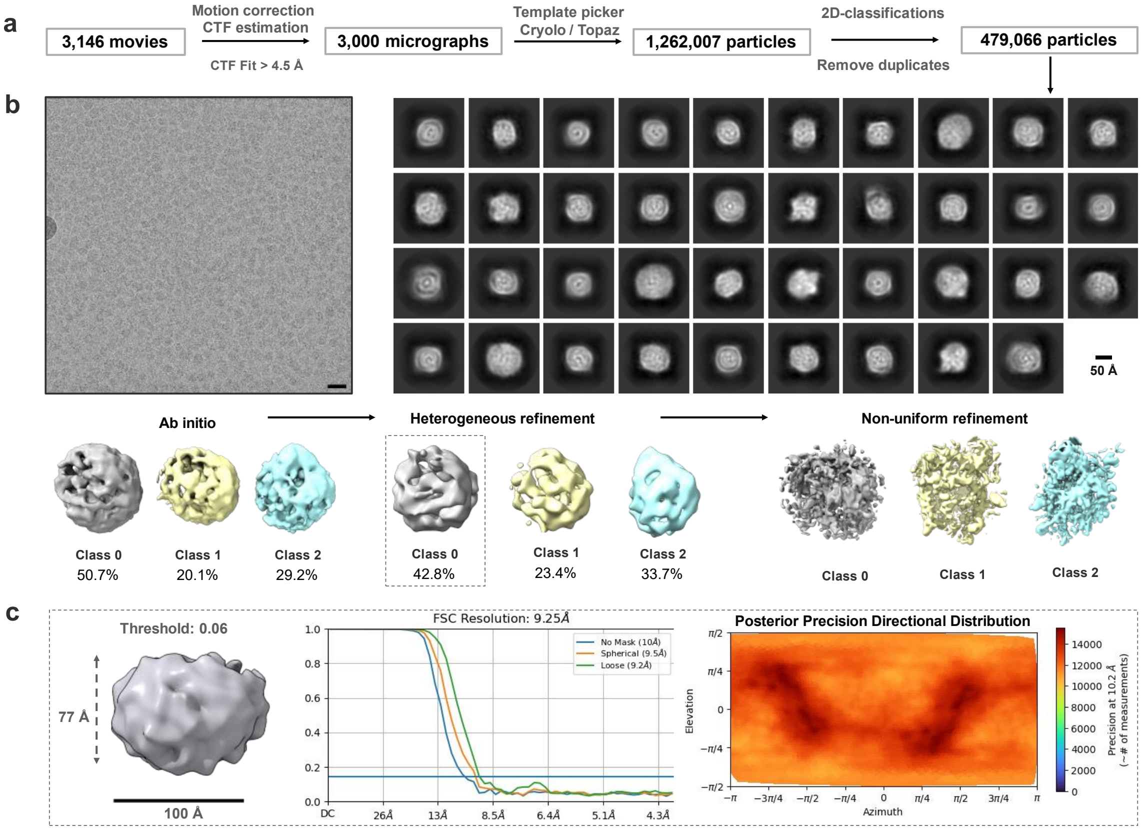


**Supplementary Fig. 4. CryoEM analysis for isNS1ts. (a)** Data analysis workflow and the corresponding number of images and particles **(b)** Representative motion-corrected micrograph with black scale bar of 20 nm, 2D classes of the picked particles, and the 3D classes generated from ab initio to refinement stages, with the particle distribution as labelled. Scale bar for 2D classes, 50 Å, as labelled in the figure. **(c)** Dimensions of a selected 3D map boxed out in panel b, black bar 100 Å, with its corresponding FSC resolution chart and the heat map of the posterior precision directional distribution as generated in crysosparc4.0[8].

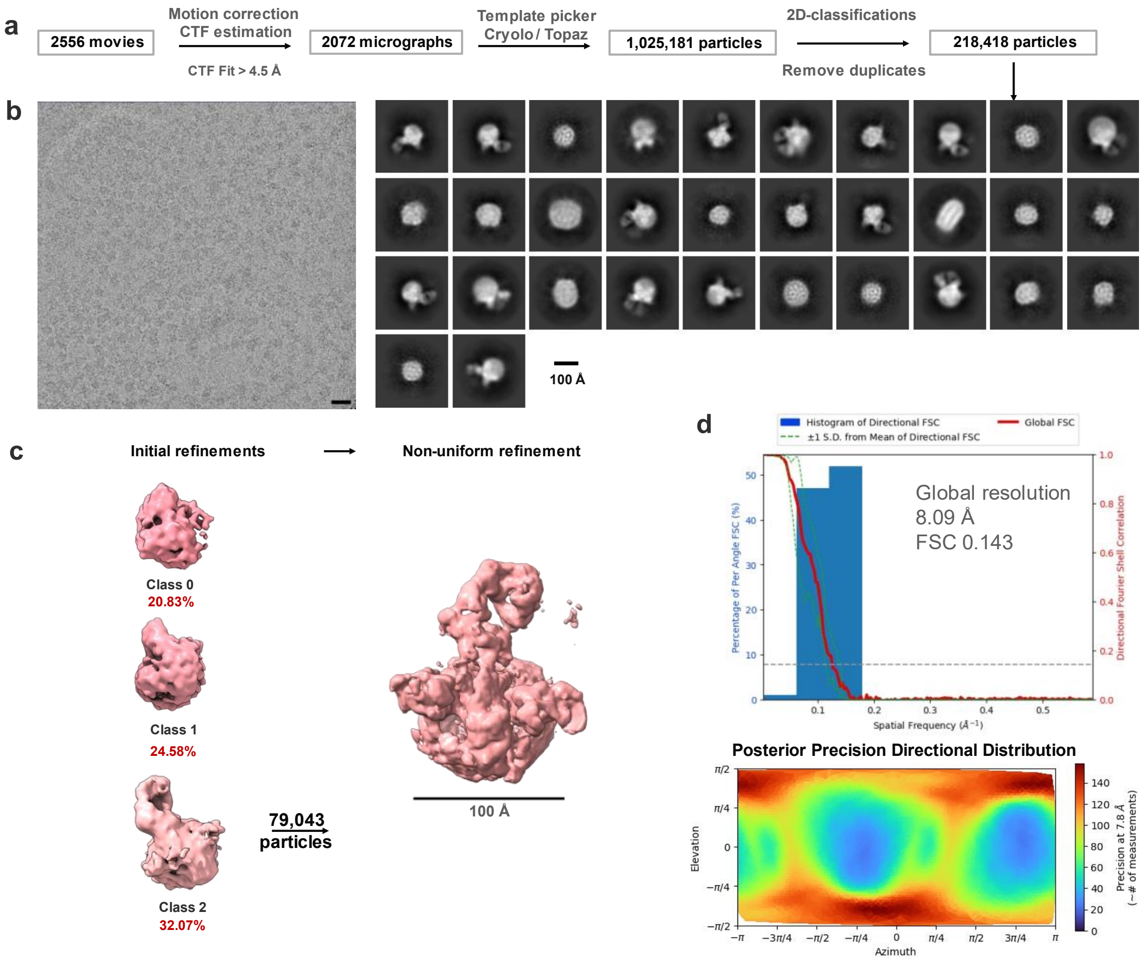


**Supplementary Fig. 5. CryoEM analysis for isNS1wt in complex with Ab56.2. (a)** Data analysis workflow and the corresponding number of images and particles. **(b)** Representative motion-corrected micrograph with black scale bar of 20 nm, 2D classes of the picked particles. Scale bar for 2D classes, 100 Å, as labelled in the figure. **(c)** Representative 3D classes contoured at 0.06 and its particle distribution as labelled and coloured in cyan. Scale bar of 100 Å as shown. **(d)** Histogram and directional FSC plot with a sphericity of 0.836 out of 1.0 generated using the remote 3DFSC processing server[14]. The global FSC (red curve) overlaid onto a histogram plot computed from all individual angular FSC values corresponding to +/−1 standard deviation of the FSC values within the histogram (green dashed curves). The corresponding posterior precision directional distribution as generated in crysosparc4.0[8].

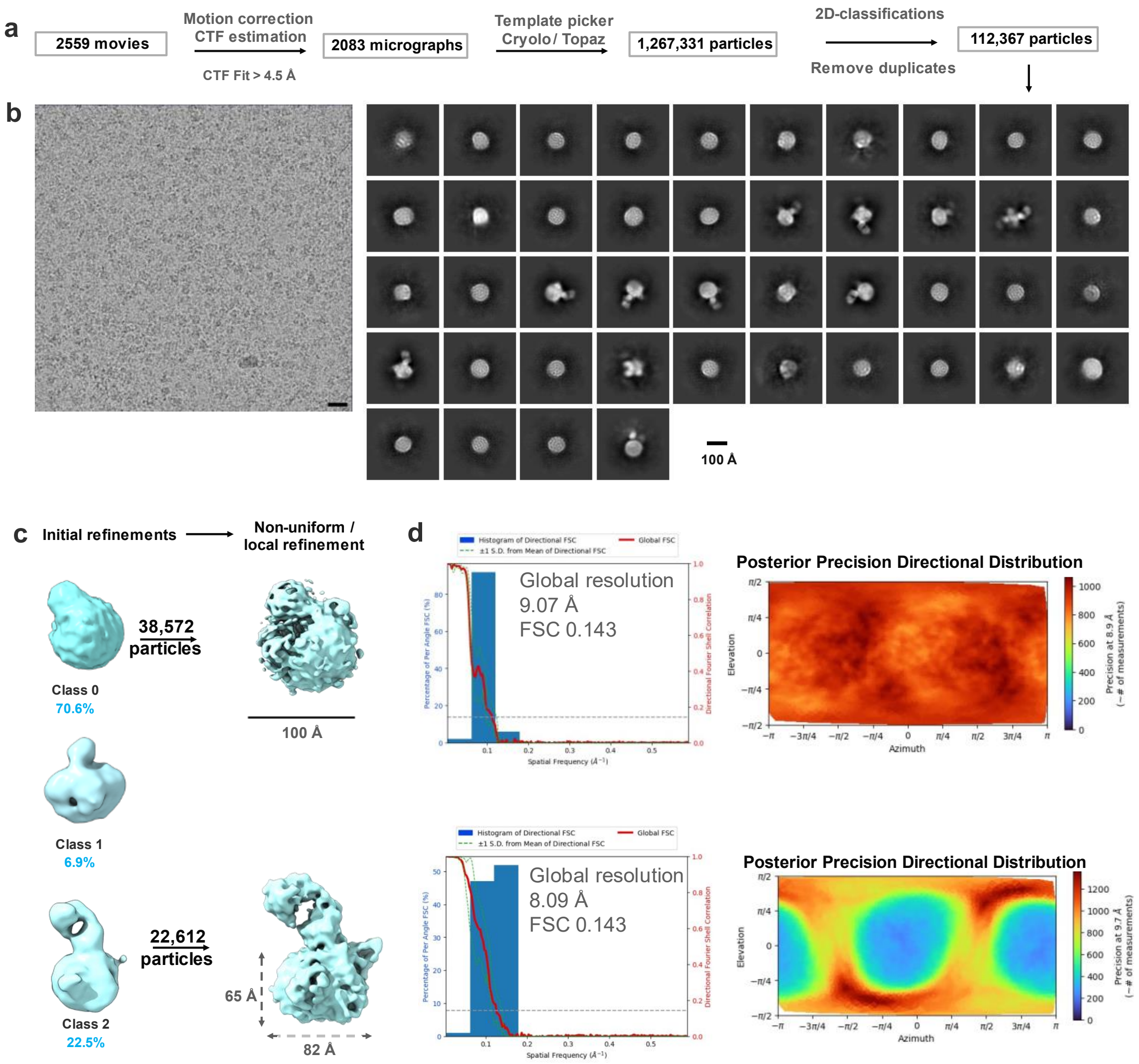


**Supplementary Fig. 6. CryoEM analysis for infection-derived isNS1wt:Fab56.2. (a)** Data analysis workflow and the corresponding number of images and particles **(b)** Representative motion-corrected micrograph with black scale bar of 20 nm, 2D classes of the overall picked particles, and initial 3D density maps generated from ab initio with the particle distribution as labelled. Scale bar for 2D classes and 3D map, 100 Å, as labelled in the figure. **(c)** 3D map classes generated from ab initio to refinement stages with the particle distribution as labelled. Refined 3D map for Class 2 contoured at 0.1, scale bar of 100 Å. **(d)** Histogram and directional FSC plot with a sphericity of 0.836 out of 1.0 generated using the remote 3DFSC processing server[14]. The global FSC (red curve) overlaid onto a histogram plot computed from all individual angular FSC values corresponding to +/−1 standard deviation of the FSC values within the histogram (green dashed curves). The corresponding posterior precision directional distribution as generated in crysosparc4.0[8].

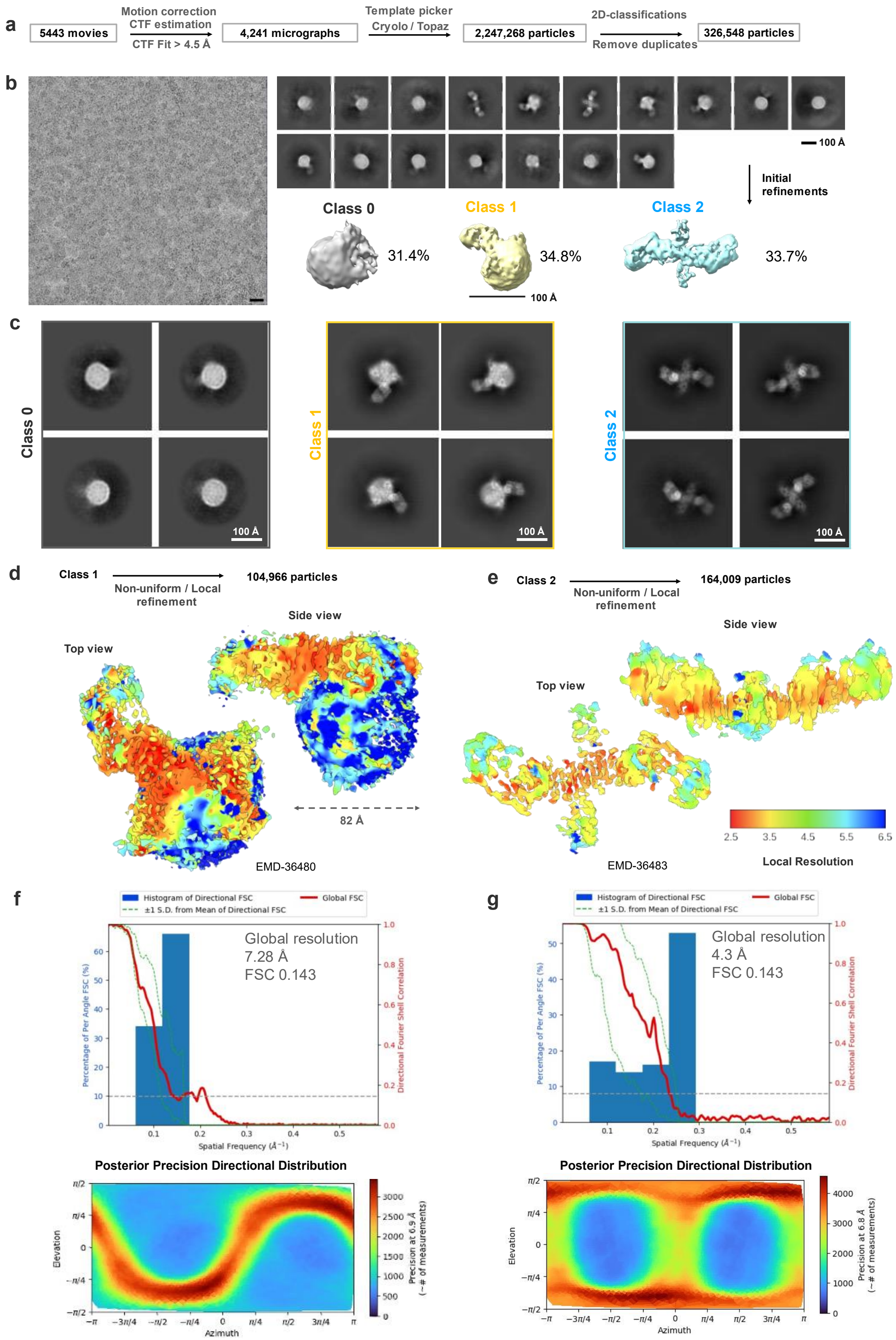
a
5443 movies
Motion correction
CTF estimation
CTF Fit > 4.5 Å
4,241 micrographs
Template picker
Cryolo / Topaz
2,247,268 particles
2D-classifications
Remove duplicates
326,548 particles
b
100 Å
Initial refinements
Class 0
31.4%
Class 1
34.8%
Class 2
33.7%
100 Å
c
Class 0
Class 1
Class 2
100 Å
d
Class 1
Non-uniform / Local refinement
104,966 particles
Top view
Side view
82 Å
EMD-36480
e
Class 2
Non-uniform / Local refinement
164,009 particles
Top view
Side view
EMD-36483
2.5 3.5 4.5 5.5 6.5
Local Resolution
f
Histogram of Directional FSC
±1 S.D. from Mean of Directional FSC
Global FSC
Global resolution
7.28 Å
FSC 0.143
Percentage of Per Angle FSC (%)
Directional Fourier Shell Correlation
Spatial Frequency (Å⁻¹)
Posterior Precision Directional Distribution
Elevation
Azimuth
Precision at 6.9 Å
(~# of measurements)
g
Histogram of Directional FSC
±1 S.D. from Mean of Directional FSC
Global FSC
Global resolution
4.3 Å
FSC 0.143
Percentage of Per Angle FSC (%)
Directional Fourier Shell Correlation
Spatial Frequency (Å⁻¹)
Posterior Precision Directional Distribution
Elevation
Azimuth
Precision at 6.8 Å
(~# of measurements)

**Supplementary Fig. 7. CryoEM analysis for isNS1ts in complex with Fab56.2. (a)** Data analysis workflow and the corresponding number of images and particles **(b)** Representative motion-corrected micrograph with black scale bar of 20 nm, 2D classes of the overall picked particles, and initial 3D density maps generated from ab initio with the particle distribution as labelled. Scale bar for 2D classes and 3D map, 100 Å, as labelled in the figure. **(c)** Representative 2D classes of each 3D classes, white bar 100 Å. Refined 3D map for **(d)** class 1 (Fab56.2:isNS1ts:HDL; EMDB: 36480) and **(e)** class 2 (Fab56.2:isNS1ts; EMDB: 36483) are coloured by local resolution and contoured at 0.1 and 0.3 respectively. The respective FSC resolution charts and the heat maps of the posterior precision directional distribution were as generated in crysosparc4.0. Histogram and directional FSC plot for **(f)** class 1 and **(g)** class 2 with a sphericity of 0.848 and 0.802 out of 1.0 generated using the remote 3DFSC processing server[14] respectively. The global FSC (red curve) overlaid onto a histogram plot computed from all individual angular FSC values corresponding to +/−1 standard deviation of the FSC values within the histogram (green dashed curves). The corresponding posterior precision directional distribution for class 1 (f, bottom) and class 2 (g, bottom) as generated in crysosparc4.0[8].

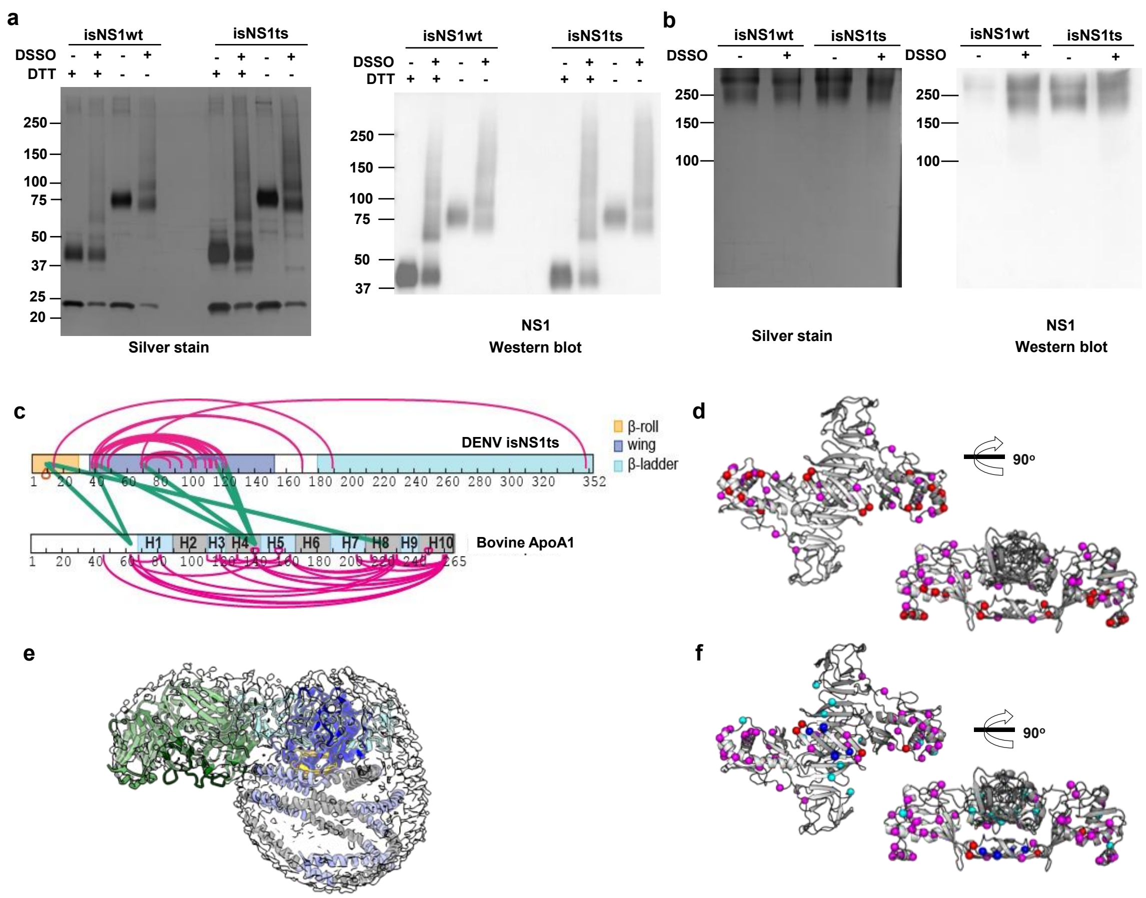


**Supplementary Fig. 8. Crosslinking mass spectrometry of sNS1:ApoA1 complex.** 2.4 µM of total protein in the purified isNS1wt1complex and 5.8 µM of total protein in the purified isNS1ts complex were crosslinked with DSSO in an isNS1: DSSO molar ratio of 1:100 for 45 minutes at room temperature. The crosslinking reaction was then quenched with 20 mM Tris-HCl (pH 8.0) for 15 minutes at room temperature. Uncrosslinked and crosslinked isNS1wt and isNS1ts were ran under reducing (with DTT) and non-reducing (without DTT) conditions in a SDS-PAGE **(a)** as well as under Native conditions in a Native-PAGE **(b)**. The gels were subjected to both silver stain and a NS1 western blot to validate NS1 bands seen in the silver-stained gels. **(c)** The residue map of sNS1ts and ApoA1 showing the intra- (magenta) and inter- (green) molecular crosslinks. The sNS1 β–roll, wing and β–ladder domains are colored in orange, blue and cyan. The repeated alpha helices of ApoA1 are colored in alternating gray and light purple. **(d)** Inter- and intra-molecular crosslinks identified for NS1ts:ApoA1 complex. The residues involved in the intramolecular crosslinks are depicted as magenta spheres and those in intermolecular cross-links are in red. **(e)** Rigid-body fitting of the overall model into the Fab-sNS1ts-HDL density map in grey and contoured at 0.17. **(f)** Superimposition between sNS1wt and sNS1ts intra- and inter-molecular crosslinks. sNS1ts inter- and intra- molecular crosslinks are depicted as magenta and red spheres while the sNS1wt intra- and inter-molecular crosslinks are as cyan and blue spheres, respectively.

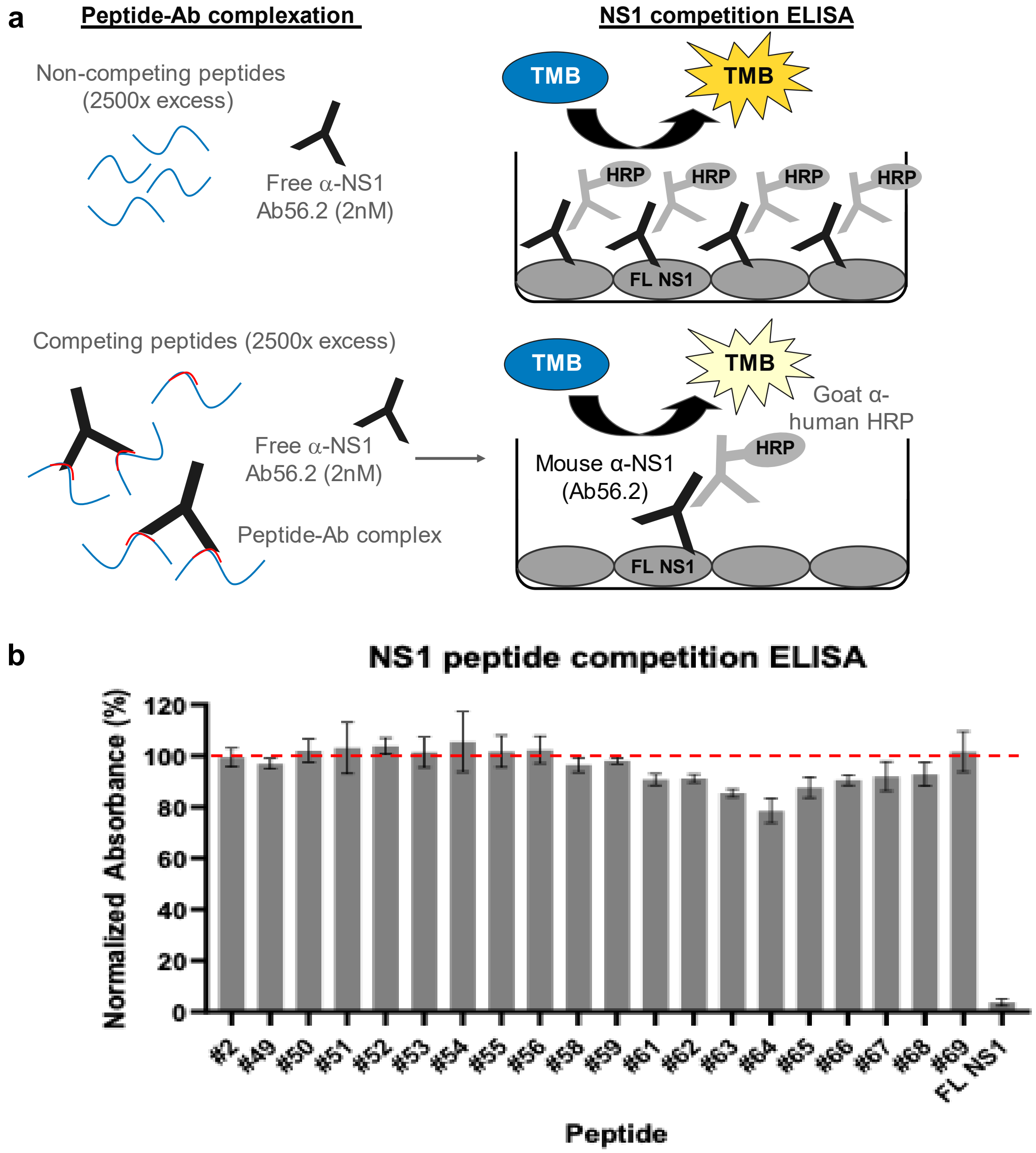


**Supplementary Fig. 9. Epitope mapping of Ab56.2 by NS1 peptide competition ELISA. (a)** Schematic of the principle of NS1 peptide competition ELISA to determine the epitope of Ab56.2. NS1 peptides (15-mer, 2500× molar excess) were first incubated with 2 nM of Ab56.2 at room temperature for formation of peptide-Ab complex. The complexation mixture is then added to ELISA plate pre-coated with full length (FL) DENV2 NS1. If the peptides contain epitope that is recognized by Ab56.2, the peptides will compete with the FL NS1 binding to the Ab56.2 in the ELISA plate, resulting in a reduced signal compared to non-competing peptides. **(b)** Normalized absorbance (%) of Ab56.2 to the FL NS1 in the presence of NS1 15-mer peptides. The absorbance reading is normalized to the signal in the absence of the peptides. FL NS1 protein is used as a positive control for the NS1 peptide competition ELISA assay. Results shown are mean ± SD from two independent experiments. Residue sequences are listed in Supplementary Table 2. β-roll residues 6-20 (#2) was used as a non-competing peptide and the red-dashed line represents the threshold for determining the competing peptides.

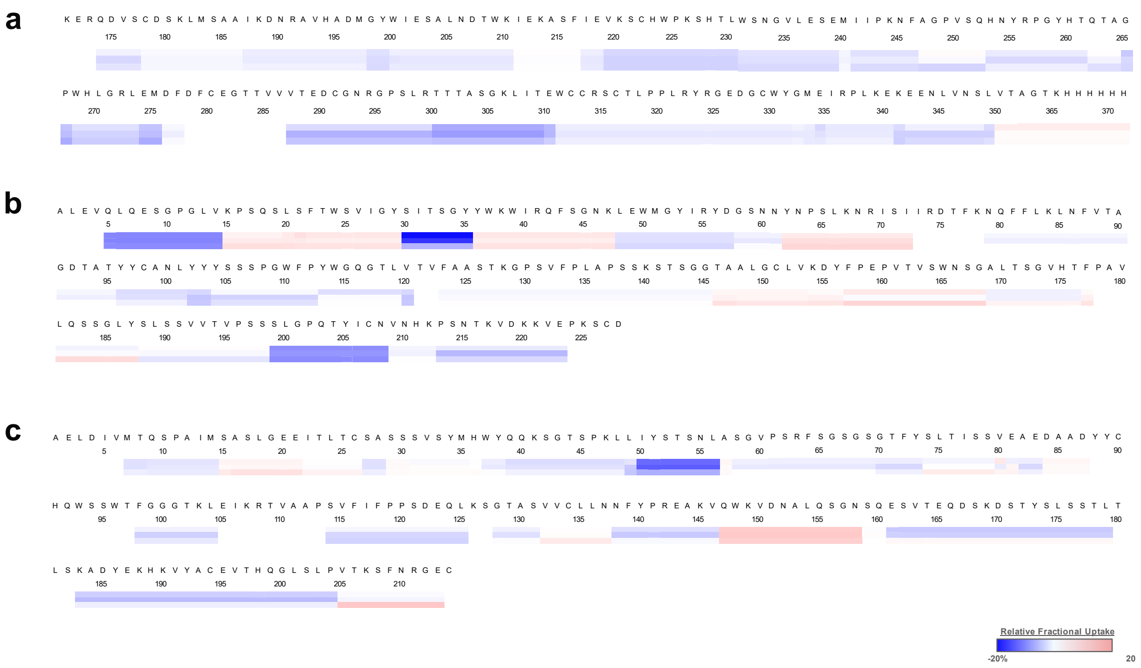


**Supplementary Figure 10.** Heat map identified specific residues associated with rsNS1c-Fab562 complex. Heat maps showing differences in relative fractional deuterium uptake for **(a)** rsNS1c, **(b)** heavy, and **(c)** light chains of Fab562. Differences were estimated by considering the deuterium exchange values of overlapping peptides across different clusters within the individual proteins of NS1c-Fab562 complex. Relative fractional uptake at 1, 10, and 100 min labeling timepoints are shown as individual lines below each residue. Heat map was generated using DynamX 3.0 software.

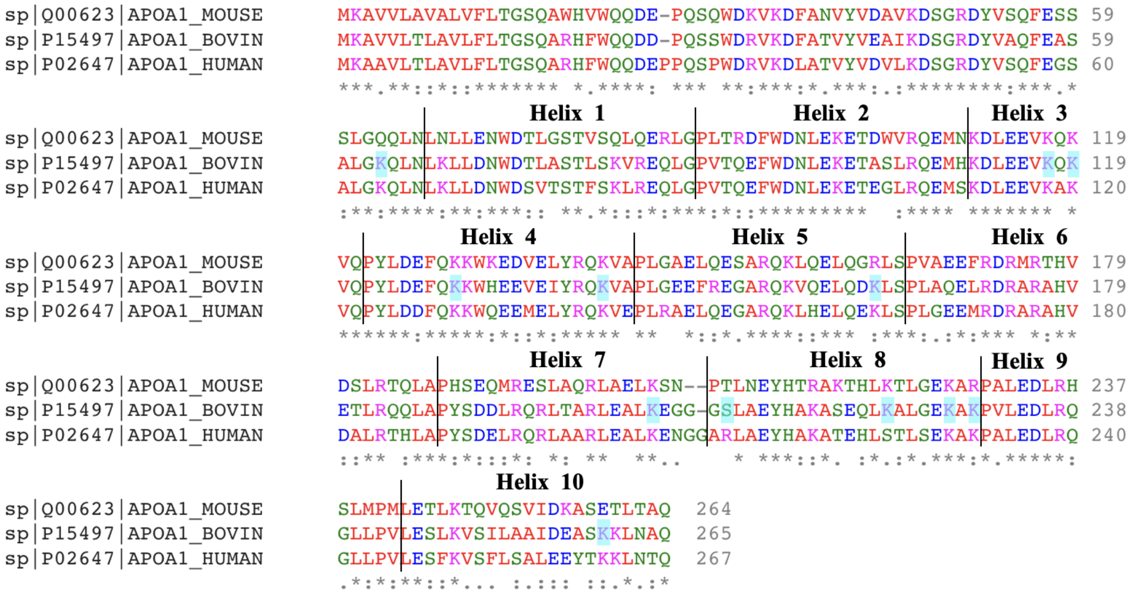


**Supplementary Fig. 11. Pairwise sequence alignment of bovine and human ApoA1**. The different helices of ApoA1 are as annotated above the aligned sequences. The residues highlighted in cyan are the crosslinked residues. The asterisk (*) signifies positions which have a fully conserved residue, the colon (:) signifies conservation between groups of strong similar properties (scoring >0.5 in the Gonnet PAM 250 matrix) and the period (.) signifies conservation between groups of weakly similar properties (scoring <0.5 in the Gonnet PAM 250 matrix) while a blank signifies dissimilar residues. Crosslinked residues identified from XL-MS are highlighted in cyan. There is a sequence similarity of 78.65% between bovine and human ApoA1, 65.17% between mouse and human ApoA1 and 66.79% between bovine and mouse ApoA1 (calculated by multiple sequence alignment, NCBI protein blast). Generated by Clustal Omega v1.2.4.

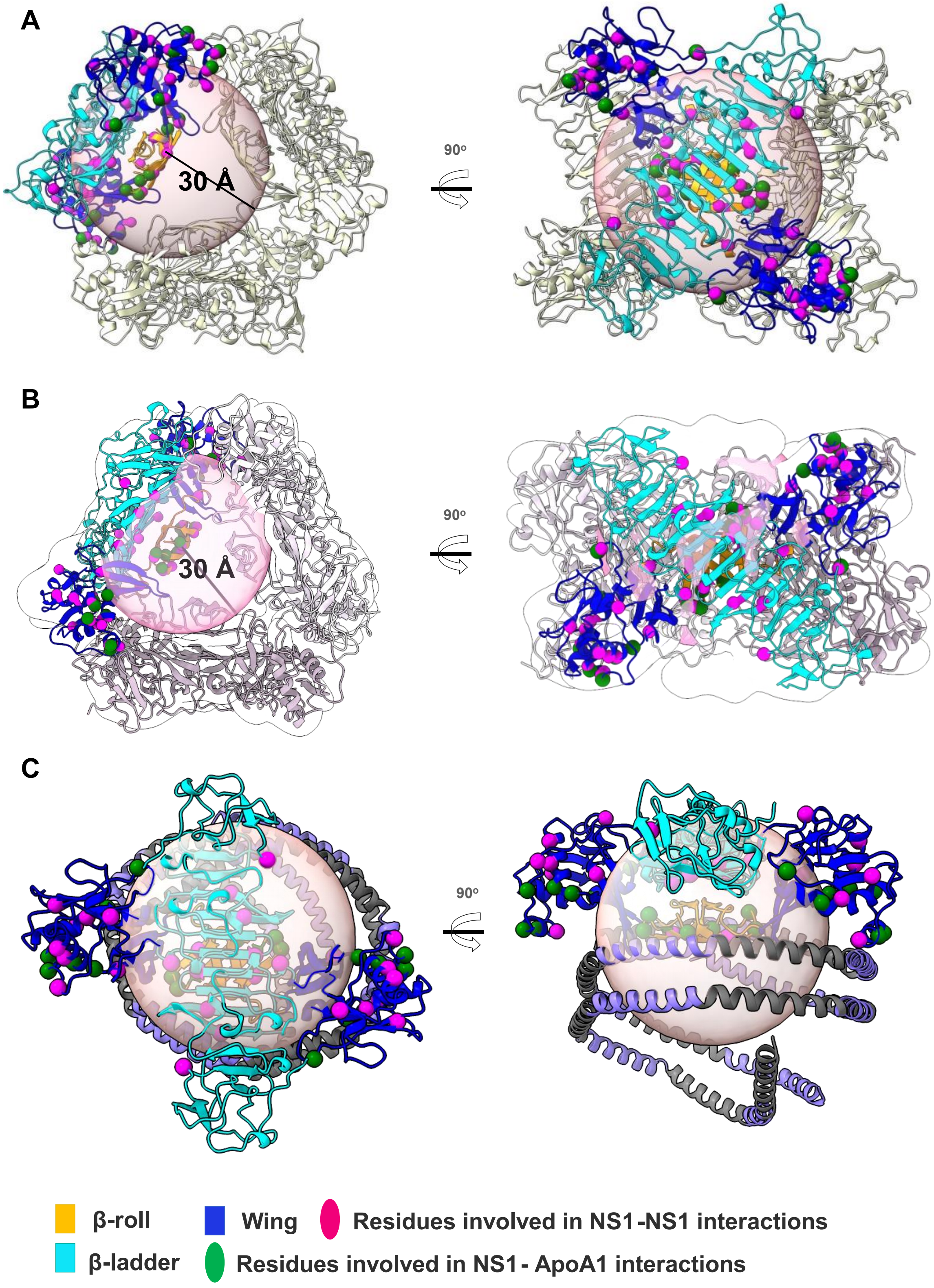


**Supplementary Fig. 12. NS1 hexamer and dimer-HDL models with residues identified in intra- and inter-molecular crosslinking interactions. (a)** Cartoon representation of isNS1hexamer due to crystal packing (PDB:4O6B) **(b)** Cartoon representation of isNS1 hexamer cryoEM model with white transparent map and black outline (EMDB:32843). The intra- and inter-molecular crosslinked residues are in magenta and green spheres. The light orange sphere with 30 Å radius represents the maximum range of crosslink-able Cα atoms from Ser2 from β-roll. **(c)** Cartoon representation of sNS1 dimer–ApoA1 cryoEM model from this study. For all panels, the top view (left panel) and side view (right panel) with intra- and inter-molecular crosslinks residues in magenta and green spheres respectively on one dimer of sNS1. Domains from the sNS1 dimer molecule are colored in orange (β-roll), blue (wing) and cyan (β-ladder). The light orange sphere has a radius of 30 Å and represent the maximum range between the Cα atoms that can be crosslinked to Ser2 residue from β-roll.

**Supplementary Table 1. Cryo-EM data collection statistics**

| | isNS1ts | isNS1wt:Ab56.2 | isNS1wt:Fab56.2 | isNS1ts:Fab56.2 |
|---|---|---|---|---|
| Grid type | gAU | gAU | gAU | gAU |
| Microscope | Titan Krios G3 | | | |
| Voltage (keV) | 300 | | | |
| Camera | Gatan K2 | Gatan K2 | Gatan K2 | Gatan K3 |
| Magnification (nominal) | 165,000 | 165,000 | 165,000 | 105,000 |
| Pixel size (Å/pixel) | 0.85 | 0.85 | 0.85 | 0.858 |
| Total electron dose (e-/Å$^2$) | 53 | 53 | 70 | 60 |
| Exposure rate (e-/Å$^2$/s) | 8.8 | 8.8 | 7 | 10.2 |
| Number of frames | 40 | 40 | 50 | 50 |
| Defocus range (µm) | 1.2-2.0 | 1.2-2.0 | 1.0-1.8 | 0.7-1.5 |
| Automation software | EPU | | | SerialEM |
| Energy filter slit width | 20 eV | | | |
| Micrographs collected (no.) | 3146 | 2556 | 2559 | 5443 |
| Micrographs post-clean (no.) | 3000 | 2072 | 2090 | 4241 |

**Supplementary Table 2.** EDEN2 NS1 peptides used in epitope mapping. Residues in red are the difference compared to D2 16681. Peptides 57 and 60 (aa residues 281-295 and 296-310 respectively) failed the QC check, no peptide made.

| Peptide # | AA Residues # | AA Residues |
|---|---|---|
| 2 | 6-20 | VSWKNKELKCGSGIF |
| 49 | 241-255 | MIIPKNFAGPVSQHN |
| 50 | 246-260 | NFAGPVSQHNYRPGY |
| 51 | 251-265 | VSQHNYRPGYHTQTA |
| 52 | 256-270 | YRPGYHTQTAGPWHL |
| 53 | 261-275 | HTQTAGPWHLGRLEM |
| 54 | 266-280 | GPWHLGRLEMDFDFC |
| 55 | 271-285 | GRLEMDFDFCEGTTV |
| 56 | 276-290 | DFDFCEGTTVVVTED |
| 58 | 286-300 | VVTEDCGNRGPSLRT |
| 59 | 291-305 | CGNRGPSLRTTTASG |
| 61 | 301-315 | TTASGKLITEWCCRS |
| 62 | 306-320 | KLITEWCCRSCTLPP |
| 63 | 311-325 | WCCRSCTLPPLRYRG |
| 64 | 316-330 | CTLPPLRYRGEDGCW |
| 65 | 321-335 | LRYRGEDGCWYGMEI |
| 66 | 326-340 | EDGCWYGMEIRPLKE |
| 67 | 331-345 | YGMEIRPLKEKEENL |
| 68 | 336-350 | RPLKEKEENLVNSLV |
| 69 | 341-352 | KEENLVNSLVTA |
